# Extreme variation in rates of evolution in the plastid Clp protease complex

**DOI:** 10.1101/405126

**Authors:** Alissa M. Williams, Giulia Friso, Klaas J. van Wijk, Daniel B. Sloan

## Abstract

Eukaryotic cells represent an intricate collaboration between multiple genomes, even down to the level of multisubunit complexes in mitochondria and plastids. One such complex in plants is the caseinolytic protease (Clp), which plays an essential role in plastid protein turnover. The proteolytic core of Clp comprises subunits from one plastid-encoded gene (*clpP1*) and multiple nuclear genes. The *clpP1* gene is highly conserved across most green plants, but it is by far the fastest evolving plastid-encoded gene in some angiosperms. To better understand these extreme and mysterious patterns of divergence, we investigated the history of *clpP1* molecular evolution across green plants by extracting sequences from 988 published plastid genomes. We find that *clpP1* has undergone remarkably frequent bouts of accelerated sequence evolution and architectural changes (e.g., loss of introns and RNA-editing sites) within seed plants. Although *clpP1* is often assumed to be a pseudogene in such cases, multiple lines of evidence suggest that this is rarely the case. We applied comparative native gel electrophoresis of chloroplast protein complexes followed by protein mass spectrometry in two species within the angiosperm genus *Silene*, which has highly elevated and heterogeneous rates of *clpP1* evolution. We confirmed that *clpP1* is expressed as a stable protein and forms oligomeric complexes with the nuclear-encoded Clp subunits, even in one of the most divergent *Silene* species. Additionally, there is a tight correlation between amino-acid substitution rates in *clpP1* and the nuclear-encoded Clp subunits across a broad sampling of angiosperms, suggesting ongoing selection on interactions within this complex.

## Introduction

Rates of sequence evolution vary dramatically across genes and genomes. Understanding the forces that determine such variation is one of the defining goals in the field of molecular evolution. In seed plants, the plastid genome (plastome) generally evolves two-to six-fold more slowly than the nuclear genome (Wolfe *et al*., 1987; Drouin *et al*., 2008; Smith and Keeling, 2015). However, among angiosperms, there is considerable heterogeneity in the rate of plastome evolution. Many lineages have maintained a slowly-evolving plastome, while others have experienced drastic rate increases (Jansen *et al*., 2007). For instance, among close relatives within the tribe *Sileneae* there have been at least three recent and independent increases in plastome evolutionary rate (Erixon and Oxelman, 2008; Sloan, Triant, Forrester, *et al*., 2014). Similar accelerations have been documented in the Campanulaceae (Haberle *et al*., 2008; Barnard-Kubow *et al*., 2014; Knox, 2014), Geraniaceae (Guisinger *et al*., 2008; Weng *et al*., 2014), Fabaceae (Magee *et al*., 2010), and Poaceae (Guisinger *et al*., 2010), among other taxa. At a structural level, plastome gene order has largely been conserved, with most angiosperms retaining the structural organization that was present in the most recent common ancestor of this group (Raubeson and Jansen, 2005). However, the sporadic increases in rates of plastome sequence evolution have often been accompanied by structural changes, including indels, inversions, duplications, shifts in inverted-repeat boundaries, and gene and intron loss (Jansen *et al*., 2007; Guisinger *et al*., 2010; Weng *et al*., 2014; Sloan, Triant, Forrester, *et al*., 2014).

Interestingly, increases in plastome evolutionary rate are often not genome-wide; rather, these increases tend to affect a subset of genes (Guisinger *et al*., 2008; Sloan, Triant, Forrester, *et al*., 2014). Commonly affected genes include those encoding the essential chloroplast factors Ycf1 and Ycf2 (Sloan, Alverson, Wu, *et al*., 2012), RNA polymerase subunits (Blazier *et al*., 2016), ribosomal proteins (Guisinger *et al*., 2008), and the AccD subunit of the acetyl-CoA carboxylase enzyme complex (Rockenbach *et al*., 2016). Some of the most striking and extreme accelerations are found in *clpP1*, which encodes a core subunit of the plastid caseinolytic protease (Clp) (Erixon and Oxelman, 2008; Barnard-Kubow *et al*., 2014; Zhang *et al*., 2014; Williams *et al*., 2015).

The plastid Clp complex plays an important role in maintaining homeostasis in plant cells by stabilizing the plastid proteome. It is the most abundant stromal protease in developing chloroplasts and has been shown to degrade various chloroplast proteins, e.g., the cytochrome b_6_f complex, a copper transporter, glutamyl-tRNA reductase, and phytoene synthase (Majeran *et al*., 2000; Nishimura and van Wijk, 2015; Apitz *et al*., 2016; Nishimura *et al*., 2017; Welsch *et al*., 2018). Consistent with the significance of the Clp complex, plastid-encoded ClpP1 as well as most nuclear-encoded subunits are essential for plant growth and viability (Kuroda and Maliga, 2003; Clarke *et al*., 2005; Zheng *et al*., 2006; Koussevitzky *et al*., 2007; Kim *et al*., 2009; Nishimura and van Wijk, 2015).

The proteolytic core of the Clp complex is composed of two stacked heptameric rings (Yu and Houry, 2007; Nishimura and van Wijk, 2015). In most bacteria, including *E*. *coli*, these 14 subunits are identical, encoded by a single gene (*clpP*). The plastid Clp core is also composed of two heptameric rings, but there have been multiple duplications of the *clpP* gene throughout cyanobacterial and plastid evolution such that the rings now comprise numerous paralogous subunits (Peltier *et al*., 2001; Majeran *et al*., 2005; Olinares, Kim, Davis, *et al*., 2011; Nishimura and van Wijk, 2015). In green plants, only *clpP1* is retained in the plastome, and the other paralogs are found in the nucleus. In *Arabidopsis thaliana*, there are a total of eight nuclear paralogs, and subunit composition differs between the two core rings. The “R ring” contains three copies of the plastid-encoded ClpP1 subunit and one of each of four “ClpR” subunits (ClpR1-4), while the “P ring” contains four different nuclear-encoded “ClpP” subunits (ClpP3-6) in a 1:2:3:1 ratio (Olinares, Kim and van Wijk, 2011). Catalytic activity in Clp subunits is conferred by an amino-acid triad (Ser 101, His 126, Asp 176). The distinguishing feature between ClpP and ClpR subunits is that the latter have each lost at least one catalytic residue and are thus assumed to be non-proteolytic; ClpR subunits are also found in the cyanobacterium *Synechococcus elongatus* and the apicoplast of the parasite *Plasmodium falsiparum* (Schelin *et al*., 2002; Peltier *et al*., 2004; El Bakkouri *et al*., 2013). Therefore, ClpP1 (the sole plastid-encoded subunit) appears to be the only catalytic member of the R ring. In addition to the core, the plastid Clp complex also contains several chaperone, accessory, and adaptor subunits required for proper Clp function, all of which are nuclear-encoded (Nishimura and van Wijk, 2015).

The importance of the plastid Clp system to cellular function makes accelerations in *clpP1* evolutionary rate particularly surprising. Various mechanisms have been hypothesized to explain such cases of rapid plastid gene evolution (Erixon and Oxelman, 2008; Guisinger *et al*., 2008; Magee *et al*., 2010; Williams *et al*., 2015; Blazier *et al*., 2016), but the relative contributions of adaptive evolution, relaxed selection, outright pseudogenization, and increased mutation rates remain unclear. Because Clp subunits are encoded by two different genomes, cytonuclear interactions are integral to the functioning of this complex. There is growing evidence that accelerated organelle genome evolution can lead to correlated increases in rates of evolution in interacting nuclear-encoded proteins. Such effects have been detected in the plastid Clp complex (Rockenbach *et al*., 2016), as well as plastid and mitochondrial ribosomes (Sloan, Triant, Forrester, *et al*., 2014; Weng *et al*., 2016), the plastid-encoded RNA polymerase (Zhang *et al*., 2015), mitochondrial oxidative phosphorylation (OXPHOS) complexes (Havird *et al*., 2015; Li *et al*., 2017; Yan *et al*., 2018), and DNA replication and repair machinery that directly interacts with plastid and mitochondrial genomes (Zhang *et al*., 2016; Havird *et al*., 2017). In plants, however, such studies have been limited to close relatives within just two groups (Geraniaceae and *Silene*) and have not been examined at deeper timescales across angiosperms.

Here, to better understand the context and scope of plastid *clpP1* acceleration, we provide a detailed accounting of the molecular evolutionary history of *clpP1* across the entire green plant lineage. Further, we use proteomic techniques to assess the functional status of *clpP1* in two species within the genus *Silene*; this genus has some of the highest and most variable observed rates of divergence for this gene. Finally, we determine whether coevolution between plastid- and nuclear-encoded subunits of the Clp complex is a broad and repeatable pattern across angiosperm diversity.

## Results

### ClpP1 has undergone numerous and extreme accelerations in rates of amino-acid substitution across independent lineages of green plants

Massive acceleration in ClpP1 amino-acid substitution rate has occurred multiple times across green plants (**Figure 1, Figure S1**). These accelerations are most pronounced in seed plants, with striking examples found in conifers, gnetophytes, and numerous angiosperm lineages. Therefore, previous reports of divergent ClpP1 sequences in specific lineages (Erixon and Oxelman, 2008; Barnard-Kubow *et al*., 2014; Zhang *et al*., 2014; Williams *et al*., 2015) appear to be the result of an incredibly frequent occurrence of accelerations over the course of seed plant evolution.

**Figure 1:**
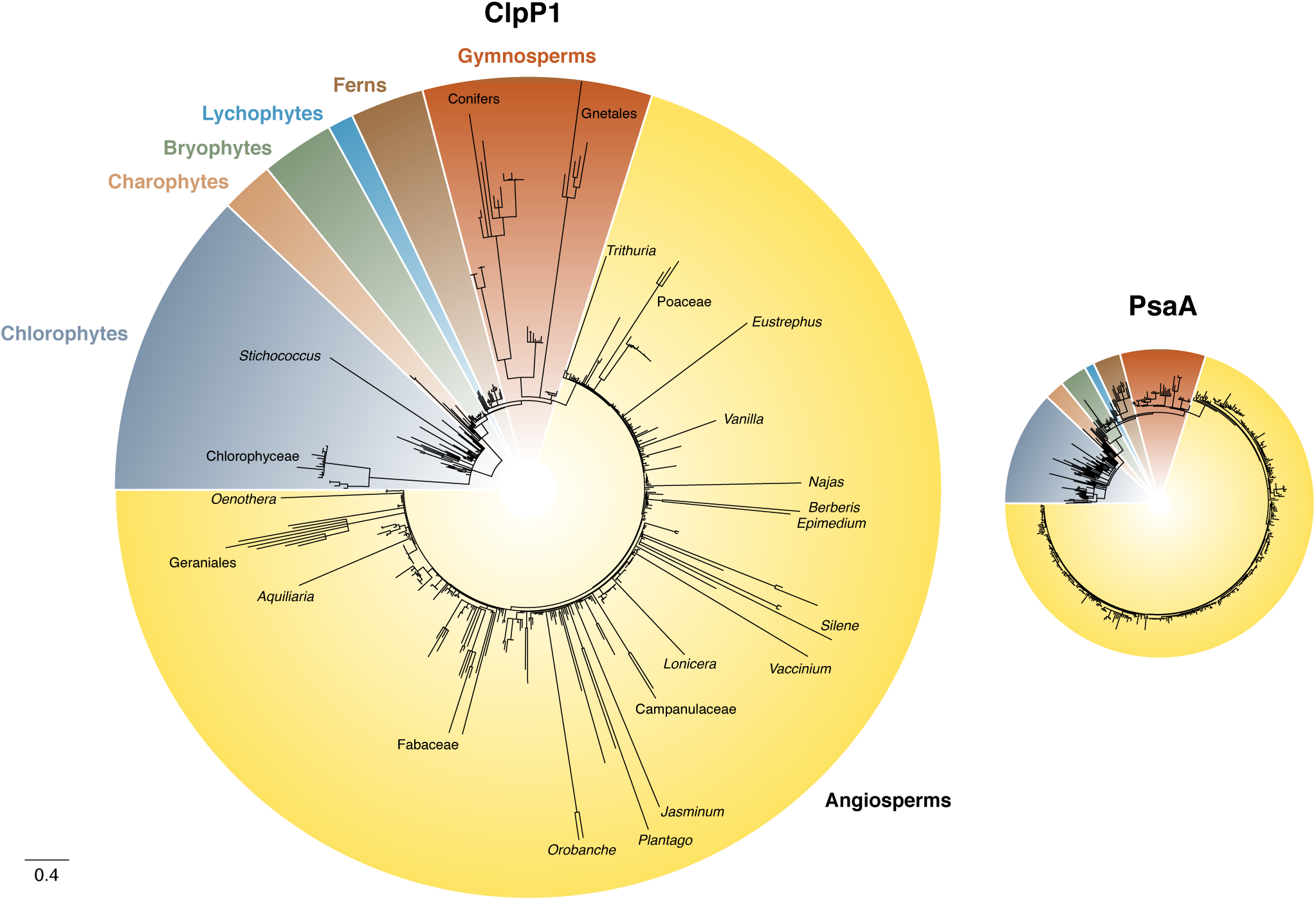
Comparison of evolutionary rates between ClpP1 and PsaA across green plants. Branch lengths represent amino acid substitutions per site. The species sampling between the trees is nearly identical (see Main Text for description of differences). Taxon names are included for select “fast” branches in the ClpP1 tree. See Figure S1 for a tree with full species labeling.

The cases of ClpP1 rate acceleration in seed plants are especially remarkable when considered against the high level of conservation in many related land plant lineages. All sampled bryophytes, lycophytes, and ferns have between 82 and 100% ClpP1 amino-acid sequence identity with the inferred common ancestor of land plants. Many seed plants show similarly high levels of conservation, but this same measure of sequence identity ranges anywhere from 27% to 87% in gymnosperms, and anywhere from 32% to 80% in angiosperms. These wide ranges correspond to major increases in amino-acid substitution rate. For instance, since the split at the base of the land plant phylogeny *ca*. 490 Mya (Morris *et al*., 2018), a representative liverwort (*Marchantia polymorpha*) and a representative hornwort (*Anthoceros angustus*) have exhibited a total rate of 0.1 amino-acid substitutions per site per billion years. At the other extreme, the angiosperms *Silene noctiflora* and *S*. *conica* diverged only about 5 million years ago (Rautenberg *et al*., 2012) and have since exhibited a rate of 335 amino acid substitutions per site per billion years – a >3000-fold increase relative to the rate of divergence between liverworts and hornworts.

To assess whether the observed rate variation in ClpP1 was the result of a plastome-wide effect, we repeated our analyses on a representative photosynthetic protein (PsaA) and found considerably less rate heterogeneity (**Figure 1**). This result is in line with the widespread conservation that is characteristic of plastid genes and especially those involved in photosynthesis (Guisinger *et al*., 2008). All sampled species of land plants, including seed plants, have between 90 and 98% identity in PsaA amino acid sequence relative to the inferred common ancestor. Further, any rate increases in PsaA have been much smaller than those in ClpP1 (Newick tree files provided at https://github.com/alissawilliams/clpP1_2018). Using the same comparisons as above between bryophytes and *Silene*, there has been a rate of 0.22 amino acid substitutions per site per billion years between *Marchantia polymorpha* and *Anthoceros angustus*, whereas this rate is 2.9 amino acid substitutions per site per billion years between *Silene noctiflora* and *S*. *conica*. Therefore, the rate has increased roughly 10-fold in *Silene* relative to the liverwort-hornwort comparison, but it is not nearly the 3000-fold increase we observe in ClpP1.

Although the most dramatic cases of ClpP1 accelerations were found in seed plants, there was also rate heterogeneity among lineages of green algae, albeit at much deeper timescales than within seed plants. The most extreme examples of ClpP1 divergence in algae are found in lineages such as *Chlorokybus, Mesostigma, Stichococcus*, and Chlorophyceae. The rapid sequence evolution within Chlorophyceae is perhaps not surprising because large insertions in ClpP1 have been previously identified in this group (Huang *et al*., 1994; Majeran *et al*., 2005; Derrien *et al*., 2012).

### ClpP1 acceleration is correlated with changes in structure and gene architecture

Our analysis demonstrated that accelerated rates of ClpP1 amino-acid substitution are also associated with broader changes at a structural level. For example, we confirmed that species with *clpP1* duplications within the plastome (**Figure S2)** generally have high rates of amino acid substitution (Erixon and Oxelman, 2008; Park *et al*., 2017). Angiosperm examples include *Silene chalcedonica* (=*Lychnis chaldeconica*), *Carex siderosticta*, and multiple lineages within the Geraniaceae. By sampling a subset of angiosperm species, we also observed that high substitution rates tend to be associated with indels in *clpP1* (**Figure S3**). This relationship was not quite statistically significant (Spearman’s rho = 0.36, p = 0.08), but that may reflect limited power resulting from our conservative approach to identifying indel events in the gappy *clpP1* sequencing alignment.

Accelerated *clpP1* evolution is also associated with intron loss in many lineages (**Figure S4**). Most land plants share two *clpP1* introns, which appear to have been gained in series during the evolution of streptophytes. Intron 1 was likely gained in a common ancestor of Charophyceae, Coleochaetophyceae, Zygnemophyceae, and land plants based on its presence in *Chara vulgaris, Chaetosphaeridium globosum*, and *Mesotaenium endlicherianum*. Intron 2 appears to have been acquired more recently in a common ancestor of Zygnemophyceae and land plants because the only algal species in which it was identified was *M*. *endlicherianum*, which is consistent with the conclusion that Zygnemophyceae is the sister lineage to all land plants (Wickett *et al*., 2014). The only other *clpP1* intron identified in our sample appears to be an independent acquisition in the streptophyte *Klebsormidium flaccidum* at a unique position within the gene. This inferred history of intron gains implies multiple secondary losses within green algae because multiple algal species within these groups lack one or both of the introns. Strikingly, we identified at least 31 independent losses of one or both introns in land plants (**Figure 2**), which are strongly associated with accelerated rates of ClpP1 evolution, including in multiple *Silene* species, the genus *Oenothera*, and *Plantago maritima*.

**Figure 2:**
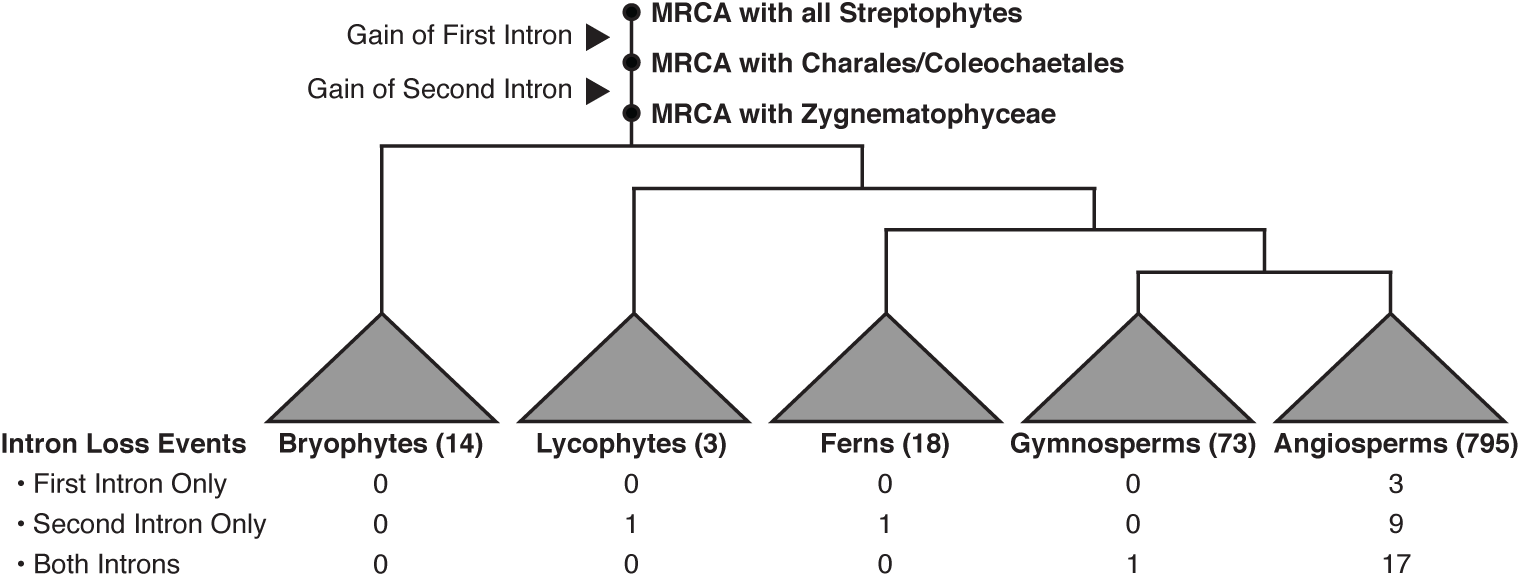
Intron gain and loss in *clpP1* across green plants. Number of species sampled is included parenthetically for each group. Columns contain the number of each type of loss in each group. MRCA, most recent common ancestor.

In land plant mitochondria and plastids, there is frequent C-to-U RNA editing (Freyer *et al*., 1997; Tsudzuki *et al*., 2001), and there is evidence that accelerated sequence evolution can also be associated with loss of editing sites (Parkinson *et al*., 2005; Sloan *et al*., 2010; Zhu *et al*., 2014; Guo *et al*., 2016). In *Arabidopsis thaliana*, there is a single RNA editing site in *clpP1* at codon 187, where the codon CAU (His) is edited to UAU (Tyr) (Tillich *et al*., 2005). This edited C has been replaced in most accelerated angiosperm species, usually in favor of “hard-coding” a T (U) at this position (**Figure S5**). In contrast, the C is maintained in most non-accelerated lineages (though there are exceptions such as the loss of this site at the base of the asterids (Hein and Knoop, 2018) well before any rate accelerations in that group). We also examined a second site in codon 28, which also undergoes CAU (His) to UAU (Tyr) editing in angiosperms (Hein and Knoop, 2018). This editing site was replaced with a hard-coded T at the base of the core eudicots but appears to be retained in other major angiosperm groups, where the same trend toward hard-coding in accelerated species is evident (**Figure S6**).

### *clpP1* loss or pseudogenization events in green plants

To assess whether the functional inactivation (i.e., pseudogenization) of *clpP1* may contribute to cases of extreme rate accelerations, we looked for evidence of potential pseudogenes and outright gene loss across green plants. Beyond the obvious loss of *clpP1* in the holoparasitic genera *Rafflesia* and *Polytomella*, which lack any detectable plastomes whatsoever (Molina *et al*., 2014; Smith and Lee, 2014), we have identified an additional seven species that appear to lack *clpP1* in their plastomes (**Table 1**). Of these seven lineages, which encompass both green algae and angiosperms, five are holoparasitic or mycoheterotrophic. The loss of photosynthesis in holoparasites and mycoheterotrophs is typically associated with radical changes in the plastome (Krause, 2008; Bromham *et al*., 2013). Although *clpP1* is often retained in these species despite massive loss of the genes that encode photosynthetic machinery (Wolfe *et al*., 1992; Delannoy *et al*., 2011), our results indicate that non-photosynthetic genes such as *clpP1* also face increased probability of loss during the evolution of holoparasitism.

**Table 1.** Examples of *clpP1* loss or putative pseudogenization

| Genus or species | Classification | *clpP1* status | Evidence |
| --- | --- | --- | --- |
| *Bathycoccus prasinos* | Green alga | Lost |  |
| *Boodlea composita* | Green alga | Lost (possibly transferred to nucleus) |  |
| *Helicosporidium sp.* | Holoparasitic green alga | Lost |  |
| *Pilostyles aethiopica* | Holoparasitic angiosperm | Lost |  |
| *Pilostyles hamiltonii* | Holoparasitic angiosperm | Lost |  |
| *Hydnora visseri* | Holoparasitic angiosperm | Lost |  |
| *Sciaphila thaidanica* | Mycoheterotrophic angiosperm | Lost |  |
| *Epipremnum aureum* | Angiosperm | Apparent pseudogenization | Large truncation/structural change |
| *Hanabusaya asiatica* | Angiosperm | Apparent pseudogenization | Loss of first exon |
| *Phelipanche purpurea* | Holoparasitic angiosperm | Apparent pseudogenization | Loss of first exon |
| *Actinidia* (except *tetramera*) | Angiosperm | Apparent pseudogenization | Loss of first exon |

\*note: *Scaevola* has also been suggested to lack *clpP1* (Jansen *et al.*, 2007), but does not have a complete plastome assembly.

In addition to cases of outright loss, there are species in which *clpP1* is still present but may be a pseudogene. For instance, the reported sequence in the angiosperm *Epipremnum aureum* is radically altered as part of a partial tandem duplication, and there are other angiosperms in which the first exon has been lost (**Table 1**). There are also lineages in which *clpP1* contains an internal stop codon. However, these cases most likely reflect posttranscriptional modifications and changes in translation rather than actual pseudogenes. The internal stop codon in the hornwort *Anthoceros angustus* has been shown to be removed by U-to-C RNA editing (Kugita *et al*., 2003), and it is possible that a similar mechanism removes internal stop codons in the hornwort *Nothoceros aenigmaticus* and the fern *Diplopterygium glaucum* – lineages in which U-to-C plastid RNA editing is prevalent (Duff and Moore, 2005; Knie *et al*., 2016). In addition, the copy of *clpP1* in the chlorophyte *Jenufa minuta* contains multiple (UGA) stop codons, but these are found at positions normally encoding conserved Trp residues in numerous genes within this plastome (Lemieux *et al*., 2015), suggesting that *J*. *minuta* has undergone a change in the plastid genetic code in which the UGA codon has been reassigned to encode Trp.

### ClpP1 is still expressed and likely assembles with nuclear-encoded Clp subunits in *Silene* species that exhibit extreme heterogeneity in rates of ClpP1 sequence evolution

To further assess whether observed increases in rates of ClpP1 sequence evolution reflect a loss of functionality, we took advantage of the rate heterogeneity within the angiosperm genus *Silene*. We selected *S*. *latifolia* as a species that has retained a highly conserved copy of ClpP1 and *S*. *noctiflora* as a close relative with a recent and extreme rate acceleration that has resulted in one of the most divergent copies of ClpP1 in our dataset, including the substitution of both histidine and aspartate in its catalytic triad (**Figure 1**). We isolated intact chloroplasts and separated the native soluble stromal complexes by native gel electrophoresis (LB-Native-PAGE), which we have applied in the past to determine the assembly state of the ClpPR core and other stromal complexes in Arabidopsis (Peltier *et al*., 2006; Olinares, Kim, Davis, *et al*., 2011). Based on tandem mass spectrometry (MS/MS) of size-fractionated samples, over 97% of all adjusted spectral counts (AdjSPC) matched annotated plastid proteins, with 1.3 and 0.9% of the AdjSPC matching to the Clp protein family in *S*. *noctiflora* and *S*. *latifolia*, respectively. We identified (i.e., deduced from the *Silene* species sequence data) all predicted ClpPRT proteins, including ClpP1 (**Figure 3; Supplemental Dataset 1**). Further, we detected ClpP1 in the same size fractions as the ClpR subunits in both species, which is consistent with the expectation of a mass of *ca*. 200 kDa for the “R-ring”(Olinares, Kim, Davis, *et al*., 2011), providing indirect evidence that ClpP1 still assembles as part of the ring structure that makes up the proteolytic core of the plastid Clp complex (**Figure 3; Figure S7)**.

**Figure 3:**
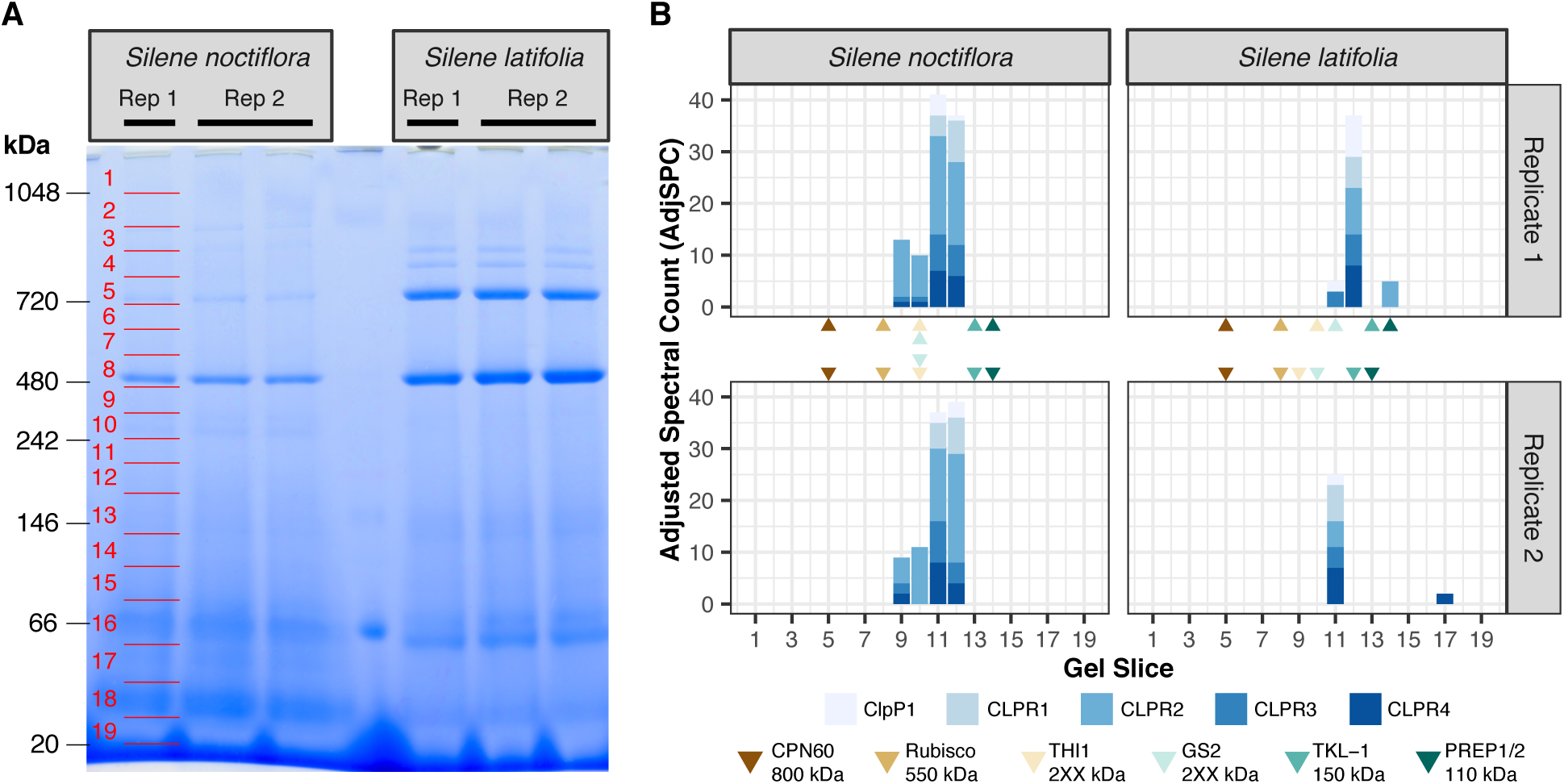
Native-gel and mass spectrometry analysis of *Silene* plastid proteins. A) LB-Native-PAGE performed on stromal protein fraction from *S*. *noctiflora* and *S*. *latifolia*. Red lines indicate approximate positions of gel slices for MS/MS analysis, but native masses were more finely calibrated with known stromal complexes. B) AdjSPC for subunits of the Clp R ring, including ClpP1. Triangles indicate the gel slice corresponding to peak detection for native complexes used for internal calibration. For more details see Figure S7 and Supplemental Dataset 1.

### Signatures of selection and rate variation across ClpP1

In species with accelerated ClpP1 sequence evolution, substitutions were widely distributed across the length of the protein (**Figure 4a**), but individual sites varied substantially in their degree of conservation (**Figure 4b**). Only a single residue (Gly at position 110) was invariant across all sampled green plants. As expected, residues in the catalytic triad were broadly conserved, though Asp 176 has been lost in over 20 species (**Figure S8)**. Substitutions at Ser 101 and His 126 were less common but still observed in 5 and 6 species, respectively. Notably, some species have experienced substitutions at multiple sites in the catalytic triad; for example, *Plantago maritima* and *Vaccinium macrocarpon* have both lost all three residues.

**Figure 4:**
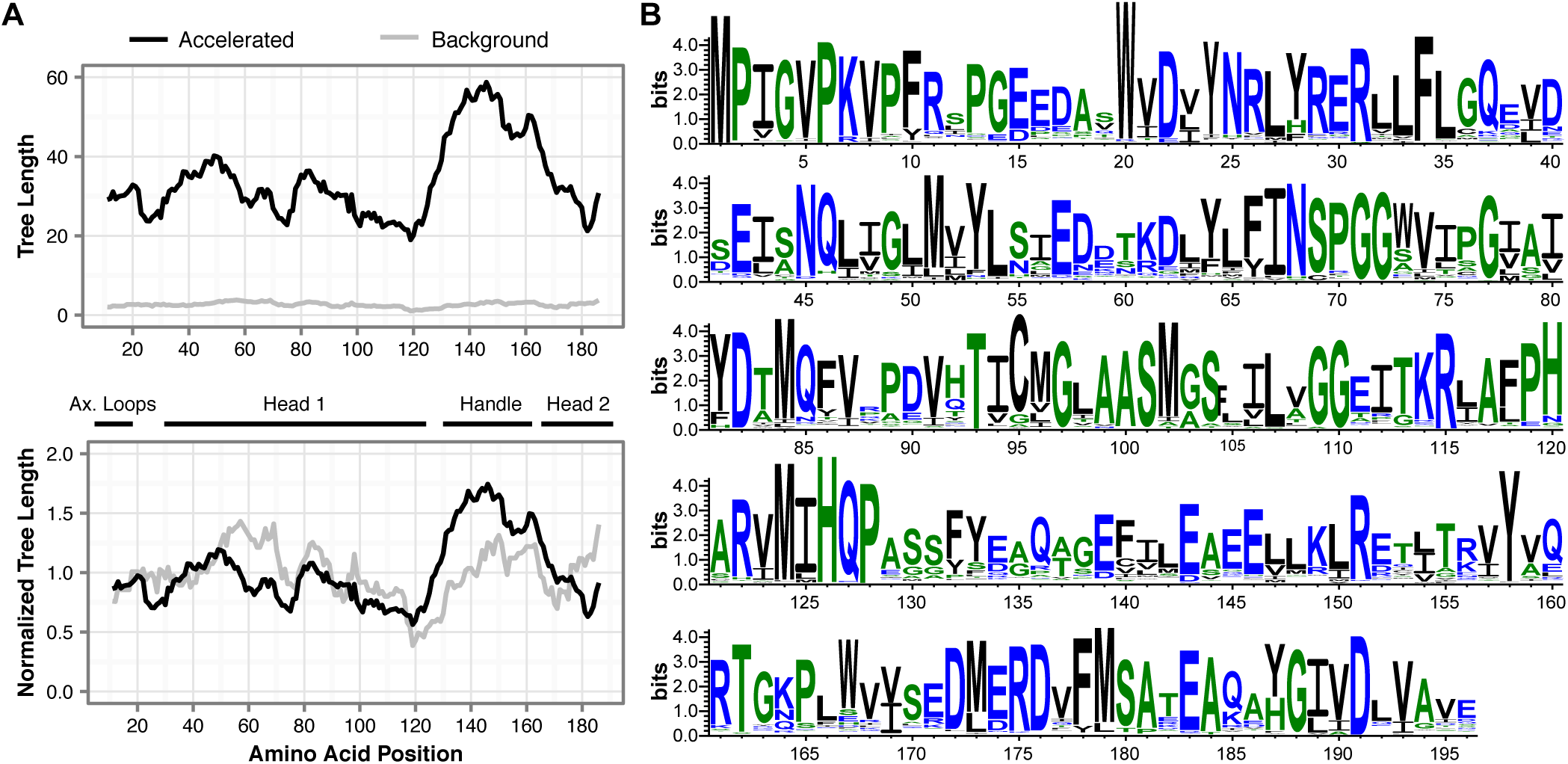
Rate variation across ClpP1. a) Sliding window analysis of rate variation across a diverse subsample of angiosperms, using a window size of 21 aa and tree length measured as amino acid substitutions per site. Normalized tree lengths (bottom plot) were calculated by dividing each window by the average tree length of the entire protein. b) WebLogo representation of sequence conservation across green plants. The size of the letter at each amino acid position is indicative of the level of conservation.

We partitioned the ClpP1 sequence into major functional domains and applied maximum-likelihood models to compare rates of amino-acid substitution across these partitions. We found that rates differed significantly among regions (χ^2^ = 337.9; d.f. = 3; *P* ≪ 0.001), with the highest rates observed in the predicted “handle” domain (**Figure 4a)**, which is likely involved in physical interactions between the two heptameric rings that make up the tetradecameric proteolytic core of the Clp complex (Yu and Houry, 2007). We also observed that the beginning of the handle domain appeared to be a hotspot for structural variants, including large insertions in *Carex, Eustrephus, Silene, Taxus*, and *Viviania* (alignment available at https://github.com/alissawilliams/clpP1_2018).

To take advantage of independent acceleration events and avoid the saturation expected when examining deep splits in the green plant phylogeny, our further analyses focused on a broad sample of 25 angiosperm species representing a wide range of rate variation. We ran a PAML branch-site model on this sample, using ClpP1-accelerated lineages as the foreground, to determine whether there are signatures of positive selection on specific codons across multiple accelerated species. This analysis identified 32 amino acid residues with greater than 95 percent probability of being under positive selection. Based on the *E*. *coli* ClpP structure, these 32 sites are scattered across the protein in no obvious spatial pattern (**Figure 5**).

**Figure 5:**
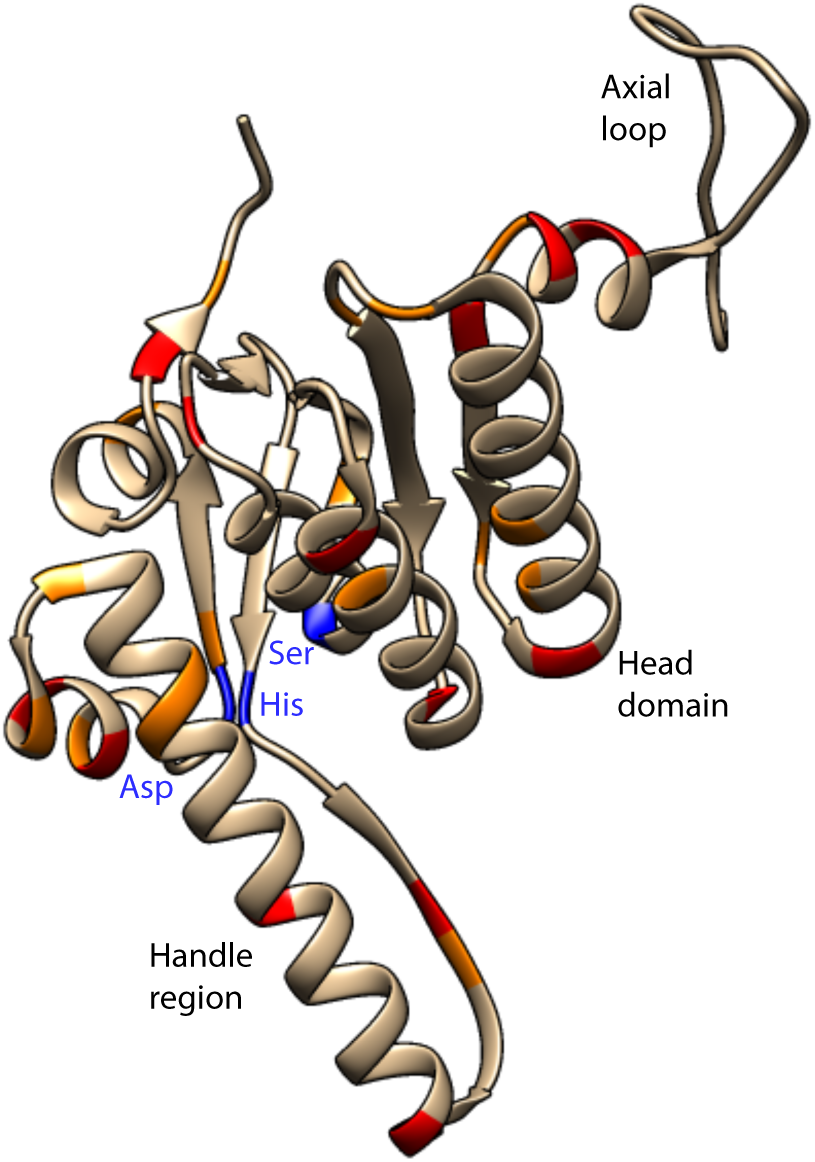
Amino acid residues under selection in ClpP1, as determined via a PAML branch-site analysis across accelerated species. Residues with posterior probabilities between 0.95 and 0.99 are colored in orange, and residues with posterior probabilities of 0.99 and above are colored in red. The three catalytic sites of ClpP1 are colored in blue. Sites were plotted on the crystal structure of the *E*. *coli* ClpP (PDB accession 1YG6).

We also calculated gene-wide average ratios of nonsynonymous to synonymous substitution rates (*d*_N_/*d*_S_) for *clpP1* and *psaA* on each branch in our angiosperm phylogeny. Across both accelerated and non-accelerated species, *clpP1 d*_N_/*d*_S_ ratios tend to be greater than the corresponding *psaA* ratios (**Figure 6**). As expected, the *clpP1 d*_N_/*d*_S_ ratios are higher in species that were identified as having highly divergent ClpP1 protein sequences. However, we found that some slow species (*Solanum lycopersicum, Cucumis sativus*, and *Vitis vinifera*) have *d*_N_/*d*_S_ ratios more characteristic of accelerated species despite much lower overall levels of ClpP1 protein sequence divergence. We obtained gene-wide *d*_N_/*d*_S_ ratios greater than 1 for *clpP1* in several species (**Table S1**) which could indicate extreme positive selection in these branches, but none of these values were significantly greater than 1 (uncorrected *p-*values all greater than 0.05). We found that variation in *d*_N_/*d*_S_ is not as large as variation in overall rates of ClpP1 protein sequence evolution, reflecting the fact that accelerated species have also experienced increased synonymous substitution rates, though not nearly to the extent of the increase in nonsynonymous substitution rates (**Figure 6**).

**Figure 6:**
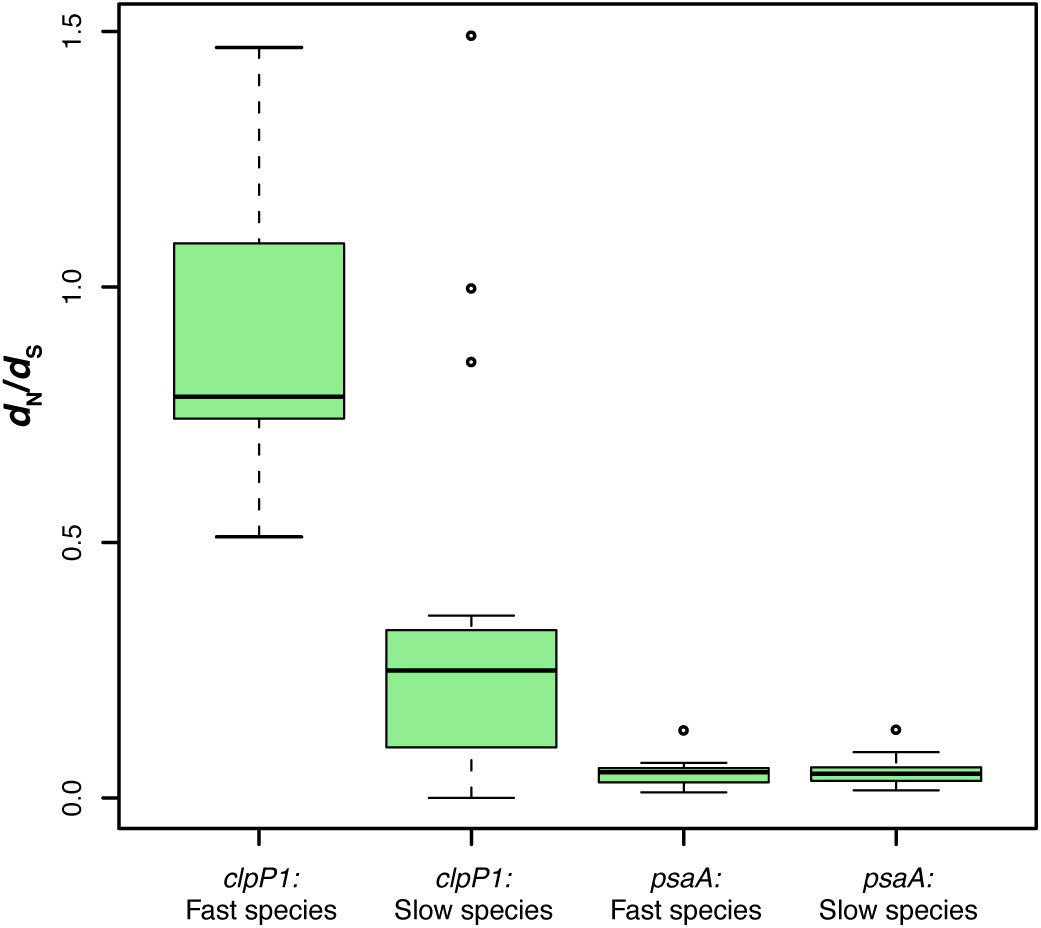
*d*_N_/*d*_S_ values for *clpP1* and *psaA* in a sample of 25 angiosperms. Species were designated as “slow” or “fast” based on ClpP1 amino acid substitution rates. The top and bottom of each box represent the upper and lower quartiles, respectively. The line contained within the box represents the median. The dotted lines connect the full range of points, apart from outliers, which are represented by dots.

### ClpP1 accelerations are highly correlated with accelerations in nuclear-encoded Clp genes

Across our sample of 25 angiosperms, the amino-acid substitution rate in ClpP1 is strongly associated with the amino acid-substitution rate of the nuclear-encoded core Clp subunits (**Figure 7a-c**). Notably, the mirroring effect between the ClpP1 and nuclear-encoded Clp branch lengths does not occur in a random sample of nuclear-encoded proteins, indicating that the correlated accelerations are not due to genome-wide rates of evolution. After using branch lengths from random genes to account for background rate variation, the vast majority of the variation in branch length for nuclear-encoded Clp subunits can be explained by the ClpP1 branch length for a particular lineage (*R*^2^ = 0.88, *P* ≪ 0.001) (**Figure 7d**).

**Figure 7:**
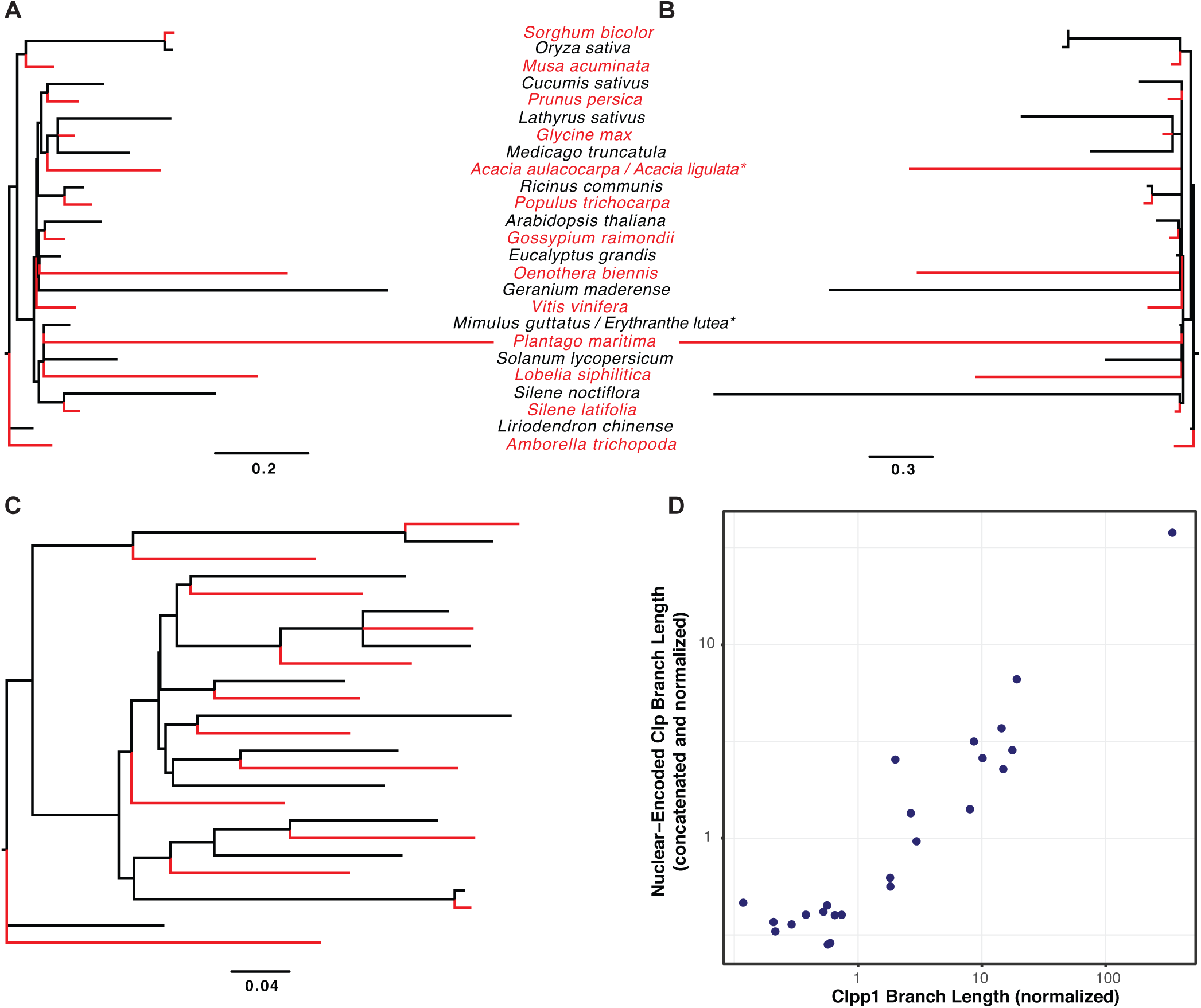
Comparison of evolutionary rates of ClpP1, the nuclear-encoded core plastid Clp subunits, and non-Clp-related nuclear-encoded genes. Species are in alternating colors for visibility. Branch lengths represent amino acid substitutions per site. a) Nuclear-encoded core plastid Clp subunits (n=8, concatenated), b) ClpP1, c) non-Clp nuclear-encoded proteins (n=20, concatenated), d) Scatterplot comparison of branch lengths. Normalization of both axes was achieved using independent sets of non-Clp nuclear-encoded proteins (n=10, concatenated).

## Discussion

### Can pseudogenization explain massive accelerations in rates of *clpP1* evolution?

Our analysis shows that accelerated *clpP1* evolution has occurred frequently and independently across green plants—particularly among seed plants. Accelerations in *clpP1* are thus a striking feature of seed plant evolution, especially given that *clpP1* is highly conserved in a majority of plant species. Pseudogenization has often been hypothesized as an explanation for extreme *clpP1* divergence (Hirao *et al*., 2008; Zhang *et al*., 2014; Williams *et al*., 2015), and in many other plastome sequencing projects, *clpP1* gene sequences have gone completely unrecognized and unannotated because of their extreme divergence (Haberle *et al*., 2008; Straub *et al*., 2011; Fajardo *et al*., 2013; Yao *et al*., 2015). There has also been speculation in these cases that *clpP1* has been functionally transferred to the nucleus, as intracellular gene transfer is a common and ongoing phenomenon in plants (Millen *et al*., 2001; Adams, Qiu, *et al*., 2002). In the highly reorganized *Boodlea composita* plastome (Cortona *et al*., 2017), *clpP1* appears to have been lost and possibly transferred to the nucleus, though we find that distinguishing *clpP1* from its nuclear paralogs is difficult in this case due to deep sequence divergence. Thus, we are not aware of any clear examples of plastid *clpP1* transfer to the nucleus in green plants.

Although we have identified probable cases of *clpP1* loss or pseudogenization within the plastome in a relatively small number of species (**Table 1**), it is unlikely that these processes represent a general explanation for the repeated and widespread pattern of substitution-rate acceleration in green plants. In the vast majority of our sampled species, *clpP1* reading frames have remained intact, even in species with extreme rates of indels and nucleotide substitutions. In the absence of functional constraints, such rapid change would quickly introduce internal stop codons and frameshifts, which suggests that there is still selection in these species to retain a functional gene copy.

Previous work has provided some evidence that even highly divergent *clpP1* genes may still be functional. For instance, the divergent copies of *clpP1* in *Acacia ligulata* and *Campanulastrum americanum* are still transcribed and spliced (Barnard-Kubow *et al*., 2014; Williams *et al*., 2015). In this study, we examined the plastid proteome of *Silene noctiflora*, a species with one of the most divergent known copies of *clpP1*, and found the first evidence that such divergent genes can still be expressed at the protein level and co-assemble with other Clp subunits. These results suggest that ClpP1 in *S*. *noctiflora* is still an important part of the core Clp structure, despite the fact that it has lost two members of its catalytic triad. If the subunit composition of R ring is indeed conserved, there are potentially important (but currently unknown) functional consequences of catalytic triad loss in the only catalytic member of this ring, which may affect overall Clp catalytic activity and/or complex structure (Andersson *et al*., 2009; Zeiler *et al*., 2013). We note that inactivation of the catalytic site in *Arabidopsis thaliana* ClpP3 has no phenotypic effect, whereas complete loss of ClpP3 results in a severe phenotype. In contrast, in case of ClpP5, loss of the catalytic site or complete gene loss results in embryo lethality (Kim *et al*., 2013; Liao *et al*., submitted).

### Role of mutation rate vs. selection in *clpP1* accelerations

Another explanation for extreme divergence in *clpP1* (and other plastid genes) could be an increase in the underlying mutation rate (Park *et al*., 2017). However, there are apparent difficulties with interpretations based solely on changes in mutation rate. While we would generally expect an increase in mutation rate to affect the entire plastome, it is clear from whole-plastome sequencing efforts that *clpP1* acceleration often occurs with little or no change in rates for a large fraction of the plastome (Guisinger *et al*., 2008; Sloan, Triant, Forrester, *et al*., 2014; Williams *et al*., 2015). The locus-specific nature of *clpP1* accelerations was supported by our analysis of *psaA* substitution rates across green plants, which were consistently low, even in species with extremely divergent *clpP1* sequences (**Figure 1**). Thus, if *clpP1* acceleration is caused by an increase in mutation rate, it would require a highly localized effect within the plastome. While such an effect is more difficult to explain than a genome-wide increase in mutation rate, “localized hypermutation” has been suggested previously as the cause of high divergence in the plastid gene *ycf4* (Magee *et al*., 2010) and may be associated with the presence of short, repetitive sequences (Stoike and Sears, 1998). There are also several documented cases of extreme variation in synonymous substitution rates across individual mitochondrial genomes, demonstrating that mutation rates do likely vary within organelle genomes in some plants (Sloan *et al*., 2009; Zhu *et al*., 2014).

One possible mechanism for localized hypermutation is “mutagenic retroprocessing” (Parkinson *et al*., 2005; Park *et al*., 2017), which occurs when a mature transcript recombines back into the genome after reverse transcription. In this scenario, accelerated substitution rates could be explained by the relatively high error rates of reverse transcriptases (Preston, 1996; Sabot and Schulman, 2006) and/or RNA polymerases (Traverse and Ochman, 2016). Retroprocessing would be expected to affect exons and introns differently. If the recombination event involves the entire gene, introns would be lost because they are not included in mature transcripts. If the recombination event involves only part of a mature transcript, that portion would necessarily be an exon, meaning that the rate acceleration would be limited to exonic regions. Both of these predictions have empirical support. Species with accelerated *clpP1* sequences often lack *clpP1* introns (**Figure S4**) (Erixon and Oxelman, 2008; Park *et al*., 2017). Among the accelerated species that do retain their *clpP1* introns, rates of sequence evolution are much higher in exons than in introns (Erixon and Oxelman, 2008; Barnard-Kubow *et al*., 2014). Despite these observations, it is not clear why *clpP1* would specifically or preferentially undergo mutagenic retroprocessing in the plastome, particularly because most plastid genes have high transcription rates and many are transcribed at higher rates than *clpP1* (Mullet, 1993; Sanitá Lima and Smith, 2017).

Another difficulty with an explanation based solely on mutation is that, if *clpP1* has only been subject to an increased mutation rate, we would not necessarily expect an increase in *d*_N_*/d*_S_. The *d*_N_*/d*_S_ statistic is typically interpreted as a measure of selection pressure, so low values are expected for genes under purifying selection, even if the mutation rate is high. Indeed, previous work has found that increased mutation pressure can even be associated with decreased *d*_N_*/d*_S_ values in genes that remain under strong purifying selection (Wolf *et al*., 2009; Havird and Sloan, 2016). In contrast, we found a trend of increased *d*_N_*/d*_S_ values in species with fast rates of *clpP1* evolution (**Figure 6**). While this result may not be surprising given that amino acid substitution rates and *d*_N_*/d*_S_ values are interconnected, it does indicate that the rates of nonsynonymous substitutions in these lineages have increased disproportionately relative to synonymous substitutions. This result suggests that *clpP1* acceleration is due, at least in part, to a change in selective pressures on protein sequence rather than simply an increase in mutation rate. This line of argument provides an alternative interpretation for the aforementioned observation that introns do not exhibit a similar degree of accelerated sequence evolution (Erixon and Oxelman, 2008; Barnard-Kubow *et al*., 2014). Importantly, any interpretation of *d*_N_*/d*_S_ ratios comes with the caveat that they can be overestimated and model-dependent, especially in cases where there are multinucleotide mutations and/or indels in the gene of interest (Li *et al*., 2009; Stoletzki and Eyre-Walker, 2011; De Maio *et al*., 2013; Venkat *et al*., 2017).

### Why might selection pressures on *clpP1* have changed?

While localized increases in mutation rate could be involved in *clpP1* accelerations, it is unlikely that mutation rates are the only contributing factor for the reasons described above. Rather, it is likely that changes in selection are involved, and increases in *d*_N_*/d*_S_ are typically caused by some combination of relaxed selection and/or positive selection. One possible explanation is that an increased mutation rate has itself altered selection pressures. This mechanism has been previously hypothesized in legumes, where the plastid gene *ycf4* has undergone increases in evolutionary rate in several species potentially as a result of localized hypermutation (Magee *et al*., 2010). Because high mutation rates can lead to an accumulation of deleterious mutations, there may be selection for affected genes to “escape” this mutation pressure by being functionally transferred to the nucleus (Blanchard and Lynch, 2000; Magee *et al*., 2010). If functional replacement does occur, the plastid-encoded gene would no longer be needed for its original function and thus experience relaxed selection. In this scenario, the decrease in functional constraint would occur due to gene transfer/replacement, which was initially driven by increased mutation pressure (Magee *et al*., 2010).

Relaxed selection can also occur without a change in underlying mutation rate. For instance, it is possible that functional constraint on the entire Clp complex could be reduced if it simply becomes less important to cellular functioning. Such a decrease in functional constraint could conceivably occur if some of the many other plastid proteases take precedence (Nishimura *et al*., 2017). Alternatively, functional constraint may be reduced specifically on *clpP1* (as opposed to the entire complex). As described above, a likely cause of relaxed selection (or outright pseudogenization) would be replacement in the Clp complex by a nuclear-transferred copy of *clpP1* or one of the existing nuclear-encoded subunits. Such transfers/replacements frequently occur for other genes in plant organelles even in the absence of increased mutational pressure (Millen *et al*., 2001; Adams, Qiu, *et al*., 2002; Adams, Daley, *et al*., 2002). However, given that we did not find evidence for widespread *clpP1* pseudogenization and/or gene replacement and that the highly divergent ClpP1 subunit in *S*. *noctiflora* still appears to be associated with other plastid Clp core subunits, it is unlikely that functional replacement of ClpP1 in the Clp complex has broadly occurred in accelerated lineages.

The other form of selection that could be involved in *clpP1* rate accelerations and increases in *d*_N_/*d*_S_ is positive selection. Under positive selection, there is selection for change, which can lead to a superficially similar pattern of accelerated protein sequence evolution as observed under relaxed selection. Previous work has found evidence of positive selection in both nuclear- and plastid-encoded Clp core subunits (Erixon and Oxelman, 2008; Rockenbach *et al*., 2016). Often, positive selection is assumed to reflect an adaptation for a novel function or a response to an environmental change. While there is no obvious shared background environment or biological feature among *clpP1*-accelerated lineages, our understanding of Clp function (including the identities of many of its target substrates) remains incomplete, so adaptive change of the whole complex is still a viable hypothesis. Further, multiple bacterial lineages including *Bacillus thuringiensis* and cyanobacteria have undergone major Clp complex reorganizations as the result of core subunit duplication and diversification, suggesting selection to deviate from the conserved ancestral functions of Clp (Fedhila *et al*., 2002; Stanne *et al*., 2007).

Positive selection could also be related to the intimate interactions between the plastid Clp subunits encoded by different genomes. Using an evolutionary rate correlation analysis, we have shown that accelerations in *clpP1* were paralleled by similar accelerations in nuclear-encoded Clp genes across a broad range of angiosperms (**Figure 7**). Our analysis represents a general class of computational methods that detect correlated rate changes between residues, genes, and/or complexes across a phylogeny as a means to identify genes that share a functional relationship and may be coevolving or at least responding to the same selection pressures (Dutheil and Galtier, 2007; Yeang and Haussler, 2007; Clark and Aquadro, 2010; Juan *et al*., 2013). Therefore, the fact that *clpP1* and the other core plastid Clp subunits had a significant rate correlation demonstrates that the plastid- and nuclear-encoded Clp subunits are subject to shared variation in selection pressures. This result could be simply due to selection acting on the complex as a whole (as described above), or it could indicate that *clpP1* and its nuclear-encoded counterparts are coevolving because of their direct interactions within the complex. Thus, coevolution between Clp subunits could be a driver of *clpP1* acceleration. A change in one subunit could introduce pressure on the other subunits to change in response—and these subsequent changes could drive further change, creating a chain reaction. Regardless of the initial trigger, this mechanism could explain both previous observations of positive selection on Clp subunits and the high correlation between their evolutionary rates. There has been speculation that such a positive feedback loop in the plastid Clp could be due to antagonistic interactions between the plastid and nuclear genomes (Rockenbach *et al*., 2016), and recent studies have implicated other plastid loci in selfish interactions with the nucleus (Bogdanova *et al*., 2015; Sobanski *et al*., 2018), but direct evidence for this or any other trigger for coevolutionary change is currently lacking.

In summary, our analysis has characterized the remarkable extent and repeatability of *clpP1* acceleration, provided evidence of retained functionality at the protein level even for one of the most extreme cases of *clpP1* divergence, and revealed an exceptionally strong, angiosperm-wide rate correlation within this complex. However, there is much work to be done, as the specific causes and mechanisms driving rapid *clpP1* evolution remain uncertain.

## Experimental Procedures

### Extraction and filtering of plastid gene sequences

All 988 complete Viridiplantae plastome sequences available in the NCBI RefSeq collection as of May 2, 2016 were downloaded from GenBank and parsed with a custom BioPerl script to extract the annotated coding sequences and number of exons for *clpP1* and *psaA*. Nucleotide sequences for species missing after this initial step were manually extracted, either via inspection of the GenBank annotation or after a tblastn v2.2.30+ (Gertz *et al*., 2006) search using the corresponding *Arabidopsis thaliana* protein sequence as a query. Extracted sequences were then screened with custom Perl scripts to identify missing or internal stop codons and identify potential annotation errors based on gene-length outliers. Corrections were made to annotations with the aid of NCBI ORFfinder (Rombel *et al*., 2002), except in cases where internal stop codons were known or inferred to be due to U-to-C RNA editing.

To reduce redundancy in the dataset, only a single sample was chosen from genera that were represented by more than one species, except in cases where substantial variation in ClpP1 sequence was observed among congeners. This down-sampling reduced the *clpP1* dataset to 480 species (and 483 sequences because we retained divergent *clpP1* copies found within the plastomes of *Carex siderosticta* and *Silene chalcedonica*). We used the same set of species for the *psaA* analysis, except that it did not include 16 holoparasitic species that have lost *psaA* along with most or all of their photosynthesis-related genes (Krause, 2008; Bromham *et al*., 2013) or the lycophyte *Selaginella moellendorffii*, which exhibits extreme levels of RNA editing (Smith, 2009), making it difficult to estimate rates of protein sequence evolution. The resulting *psaA* dataset contained 463 sequences.

In the Methods below, we will refer to the set of Viridiplantae species described above as the “large dataset” (**Table S2**). For some analyses, we used a more targeted sampling of 25 angiosperms, which we will refer to as the “small dataset” (**Table S3**). The species in the small dataset were selected to 1) span the phylogenetic diversity of angiosperms, 2) capture multiple independent accelerations in plastome evolutionary rate as well as related species with slow evolutionary rates, and 3) only include species for which nuclear genome/transcriptome resources were available (for use in subsequent analyses of cytonuclear coevolution; see below). All scripts, sequence data, and alignments are available at https://github.com/alissawilliams/clpP1_2018.

### Sequence alignment and tree construction

To assess variation in rates of ClpP1 and PsaA protein sequence evolution across green plants, the nucleotide sequences of the large dataset were translated into protein sequences in MEGA v7.0.21 (Kumar *et al*., 2016). These protein sequences were then aligned using the MAFFT v7.222 einsi option (Katoh and Standley, 2013). Constraint trees were manually constructed based on established phylogenetic relationships, using NCBI taxonomy and the Angiosperm Phylogeny Website v13. Branch lengths were estimated using codeml in the PAML v4.9a package (Yang, 2007) with an LG substitution matrix and rate variation among sites estimated with a gamma distribution. For this analysis, the ClpP1 alignment was trimmed to remove all insertions relative to the 196-aa *Nicotiana tabacum* reference sequence.

### Analysis of substitution rate variation and tests for selection

#### Variation among sites

To generate site-specific estimates of amino-acid substitution rate across the ClpP1 subunit, we applied a partitioned model in codeml. Using option G, we specified a separate partition for each position in the 196-aa alignment. The Mgene parameter was set to 0 such that total tree length could vary across partitions (i.e., different sites could have different rates), but branch lengths had to remain proportional. The complexity of this model necessitated that it be run under a simple Poisson model of amino acid substitutions. To assess whether certain regions of the protein exhibited disproportionate accelerations in fast lineages, we performed this analysis on two different subsets of the large dataset. The first was a set of 27 slow “background” lineages sampled from across green plants. The second was a set of 60 angiosperms that contained 38 “accelerated” lineages and 22 interspersed slow lineages that were included to increase the probability of detecting parallel amino-acid substitutions in fast lineages. Site-specific rate estimates were summarized using a sliding-window analysis with a window size of 21 aa. This rate analysis was performed both on raw tree lengths and on normalized rates that were scaled to the average tree length across the data set. Site-specific variation was also summarized with WebLogo v3.5.0 (Crooks *et al*., 2004), using default settings and the trimmed 196-aa alignment of 483 ClpP1 sequences described above.

To further investigate rate variation within ClpP1, we partitioned the subunit into the following four regions based on characterized structural domains in *E*. *coli* (Yu and Houry, 2007): 1) the “handle domain”, consisting of positions 129-162; 2) the “head domain”, consisting of positions and 32-124 and 165-193; 3) the N-terminal “axial loops”, consisting of positions 7-20 (although the alignment between *E*. *coli* and the plastid ClpP1 is weak in this region, so it not clear whether there is a conserved functional role); and 4) “other” spacer regions, consisting of all remaining positions in the 196-aa alignment. We repeated the codeml analysis described above but used the full 483-sequence alignment and specified these four partitions with option G. As a basis of comparison, we performed the same analysis without any partitions. To test for evidence of significant rate heterogeneity among the four regions, we performed a likelihood ratio test (LRT) that compared these two models with three degrees of freedom.

#### Variation among branches

To examine differences in *d*_N_/*d*_S_ across species, we used codeml to determine *d*_N_ and *d*_S_ (for both *clpP1* and *psaA*) for each branch of the small dataset. We converted our previously obtained ClpP1 and PsaA amino-acid alignments into codon-based nucleotide alignments. In the codeml run, we used the parameters model=1 and fix_omega=0, which together specify estimation of an individual *d*_N_/*d*_S_ value for each branch in the tree. For terminal branches with a *d*_N_/*d*_S_ estimate > 1, we assessed statistical significance by constraining each branch of interest (separately) to a *d*_N_/*d*_S_ value of 1 (model=2, fix_omega=1). We determined whether the unconstrained PAML model was a significantly better fit to the data than each constrained model by performing LRTs with one degree of freedom.

#### Branch-site models

Using the codon-based alignment of *clpP1* for the small dataset (described above), we used a codeml branch-site model (Zhang *et al*., 2005) to infer whether any amino acid sites in ClpP1 have been subject to positive selection. We partitioned the species of the small dataset into one of two categories: fast or slow rates of ClpP1 evolution. The “fast” species were specified as the foreground, which means that the analysis identified sites under positive selection in this species subset. This analysis used the parameters model=2, NSsites=2, fix_omega=1, and omega=1 for the null model, and the parameters model=2, NSsites=2, fix_omega=0, and omega=1 for the alternative model. To determine whether the alternative model was a significantly better fit to the data than the null model, we performed an LRT with one degree of freedom. We mapped sites with a Bayes empirical Bayes (BEB) posterior probability of ≥ 0.95 for positive selection based on this analysis to the homologous positions in the *E*. *coli* ClpP structure (Protein Data Bank 1YG6) visualized in Chimera v1.11.2 (Pettersen *et al*., 2004).

### Evolutionary rate covariation between the plastid- and nuclear-encoded Clp subunits

To determine whether the rate of amino-acid substitution in ClpP1 is correlated with the rate in nuclear-encoded Clp subunits across angiosperms, we compiled protein sequences of all nine core Clp subunits (ClpP1,3-6, ClpR1-4) and 20 non-Clp nuclear-encoded genes for each species in the small dataset, using a custom Python script to reciprocally blast *A*. *thaliana* protein sequences against predicted protein sequences from each of the other species. Predicted protein sequence data were collected from various sources, including sequenced genomes on Phytozome (https://phytozome.jgi.doe.gov/), transcriptomes from the 1KP project (Wickett *et al*., 2014), and various other transcriptome sequencing projects (**Table S3**). The 20 non-Clp genes represent the subset of the 50 control genes examined by Rockenbach *et al*. (Rockenbach *et al*., 2016) for which we could recover orthologs in all species of interest. Protein sequences were aligned with MAFFT (Katoh and Standley, 2013) and trimmed at the N- and C-terminal ends as needed (in cases of poor alignment) to avoid overestimation of branch lengths. In rare cases, internal trimming was required for the same reason. In two cases, nuclear sequences from one species were paired with a ClpP1 sequence from a closely related species (*Acacia aulacocarpa* was paired with *A*. *ligulata*, and *Mimulus guttatus* was paired with *Erythranthe lutea*) due to the lack of a published plastome (**Table S3**).

We estimated branch lengths both for individual proteins and for sets of concatenated amino-acid sequences using codeml with the LG substitution matrix and a gamma distribution. The concatenated sets of sequences were 1) all nuclear-encoded core Clp proteins, 2) all nuclear-encoded non-Clp proteins, and 3) two randomly divided halves of the 20 nuclear-encoded non-Clp proteins (10 proteins each). We used correlation analysis to compare the branch lengths of ClpP1 and the concatenated nuclear-encoded Clp proteins, each normalized by dividing by the branch length of one set of concatenated non-Clp proteins in that species. By using the two different halves of the dataset for normalization, we avoided introducing statistical non-independence between our variables in the correlation analysis. Only terminal branches were used in the correlation analysis, with the exception of the grasses (*Oryza sativa* and *Sorghum bicolor*). In that case, ClpP1 acceleration occurred before the split between the two species; thus, the branch leading to the grasses was used in place of the two terminal branches of those species. We log-transformed the normalized branch lengths for ClpP1 and the concatenation of nuclear-encoded Clp subunits and calculated the Pearson correlation coefficient across branches with R v3.4.1.

### Analysis of indels

To determine the relationship between rates of amino-acid substitution and rates of indels in *clpP1*, we used the codon-based alignment for the small dataset. Indels were coded using the modified complex coding option in SeqState v1.0 (Müller, 2005). The indel data were visualized in Mesquite v3.31 using “Trace Character History ⟶ Parsimony Ancestral States.” This visualization plotted indel states on each branch of the constraint tree at each indel site. Using these plots, we counted the number of novel indels on each branch of the tree. In this case, all novel indels occurred on terminal branches. Correlation analysis was performed similarly to the plastid-nuclear Clp correlation described above. Once again, the substitution-based branch lengths for ClpP1 were normalized with one concatenated set of non-Clp sequences. The branch-specific ClpP1 indel counts were normalized with the branch lengths from the other concatenated set of non-Clp sequences. We tested for a significant association between these normalized rates of ClpP1 sequence and structural evolution, using a Spearman Rank Correlation analysis in R.

### Proteome analysis of Clp core complexes in *Silene species*

#### Tissue collection and chloroplast isolation

A total of 60 g of leaf tissue was collected from mature rosettes from four *S*. *noctiflora* individuals and from ten *S*. *latifolia* individuals. The *S*. *noctiflora* individuals were derived from the BRP line previously used for mitochondrial genome sequencing (Wu *et al*., 2015), and the *S*. *latifolia* individuals were derived from the line previously used for plastid genome sequencing (Sloan, Alverson, Wu, *et al*., 2012). Plants were grown in Fafard 2SV Mix supplemented with vermiculite and perlite in the Colorado State University greenhouses with supplemental lighting on a 16/8-hr light/dark cycle and regular watering and fertilization. Chloroplasts were isolated following a protocol based on van Wijk et al. (van Wijk *et al*., 2007). In brief, rinsed leaf tissue was disrupted with a blender in 100 ml of grinding buffer for each 10 g of tissue (50 mM HEPES-KOH pH 8.0, 330 mM sorbitol, 2 mM EDTA, 5 mM ascorbic acid, 5 mM cysteine, 0.05% BSA) and filtered through two layers of Miracloth. The resulting samples were centrifuged at 1300 x g for 4 min in a fixed-angle rotor. Pellets were resuspended in wash buffer (50 mM HEPES-KOH pH 8.0, 330 mM sorbitol, 2 mM EDTA), loaded onto 40%/85% Percoll step gradients, and centrifuged at 3750 x g for 10 min in a swinging-bucket rotor. Intact chloroplasts were harvested from the interface of the step-gradient, diluted in wash buffer, and centrifuged at 1300 x g for 3 min in a fixed-angle rotor. The resulting chloroplast pellets were flash frozen in liquid nitrogen and stored at -80 °C. All isolation steps were performed at 4 °C under dim green light.

#### Isolation of stromal protein fractions and native polyacrylamide gel electrophoresis (Native PAGE)

Frozen chloroplast pellets were resuspended in HEPES 50 mM pH 8, MgCl_2_ 10 mM, glycerol 15% and protease inhibitors, and the soluble stromal proteomes were isolated from the resuspended, broken chloroplasts by centrifugation at 100,000 x g for 30 min at 4 °C. The supernatant containing the chloroplast soluble proteomes were concentrated by Amicon 10 kDa filter units and proteins were quantified by BCA Protein Assay Kit (Thermo Fisher). For light-blue native PAGE analysis, 50 µg of stromal protein of each species in 50 mM BisTris-HCl, 50 mM NaCl, 10% w/v glycerol and 0.001% Ponceau S (pH 7.2) was loaded per lane using the NativePage Novex gel system with precast 4 - 16% acrylamide Bis-Tris gels (Invitrogen). The upper buffer contained 0.002% Coomassie G. A total of three lanes were loaded for each species.

#### Mass spectrometry (MS) and Data analysis

Each gel lane was cut into 19 bands followed by reduction, alkylation, and in-gel digestion with trypsin as described in (Shevchenko *et al*., 2006; Friso *et al*., 2011). For replicate 1, we used one gel lane for each species, whereas we pooled two gel lanes for replicate 2 to increase protein identifications and sequence coverage. The resuspended peptide extracts were analyzed by data-dependent MS/MS using an on-line LC-LTQ-Orbitrap (Thermo Electron Corp.) with details as described in (Kim *et al*., 2015). Hence, a total of 38 MS/MS runs for each species was carried out. MS data searching against assembled databases for *S*. *noctiflora* (88,166 sequences; 20,816,406 residues) and *S*. *latifolia* (101,108 sequences; 20,447,864 residues) was done using Mascot, followed by filtering, grouping of closely related sequences based on matched MS/MS spectra and quantification based on normalized AdjSPC (NadjSPC) as in (Friso *et al*., 2011; Kim *et al*., 2015). For each species, databases were a merger of annotated proteins from organellar genomes (Sloan, Alverson, Chuckalovcak, *et al*., 2012; Sloan, Alverson, Wu, *et al*., 2012) and protein sequence predictions generated by TransDecoder (Haas *et al*., 2013) from transcriptome assemblies (Sloan, Triant, Wu, *et al*., 2014). Protein annotations are based on homology to Arabidopsis and taken from the Plant Proteome Data Base (http://ppdb.tc.cornell.edu/).

To determine the assembly state of ClpP1 and other ClpPRT subunits, native masses were calibrated with endogenous stromal complexes for which we and/or others previously determined the native mass in Arabidopsis, in particular CPN60 (800 kDa), RUBISCO holocomplex (550 kDa), glutamate-ammonia ligase (GS2; 240 kDa), CLPC/D (200 kDa), transketolase-1 (TKL-1; 150 kDa), thiazole biosynthetic enzyme 1 (THI1; 245 kDa), and metalloprotease PREP1/2 (110 kDa) (see (Peltier *et al*., 2006) and references therein).

## Acknowledgements

We thank Kate Rockenbach for assistance with implementing chloroplast isolation techniques in *Silene*. We thank Dr. Nazmul Bhuiyan for native gel analysis of stromal proteomes and Dr. Qi Sun from Computational Biology Service Unit (CBSU) at Cornell University for his support with the *S*. *noctiflora* and *S*. *latifolia* databases, database searches, and protein annotations. We thank Andrea Del Cortona for providing *Boodlea composita* sequence data. This work was supported by a National Science Foundation (NSF) Graduate Research Fellowship (DGE-1321845 to A.M.W.), NSF MCB-1733227 to D.B.S., and NSF MCB-1614629 to K.J.V.W. A.M.W. was also supported by a Department of Education GAANN Fellowship.

## Supplementary Materials

**Figure S1:**
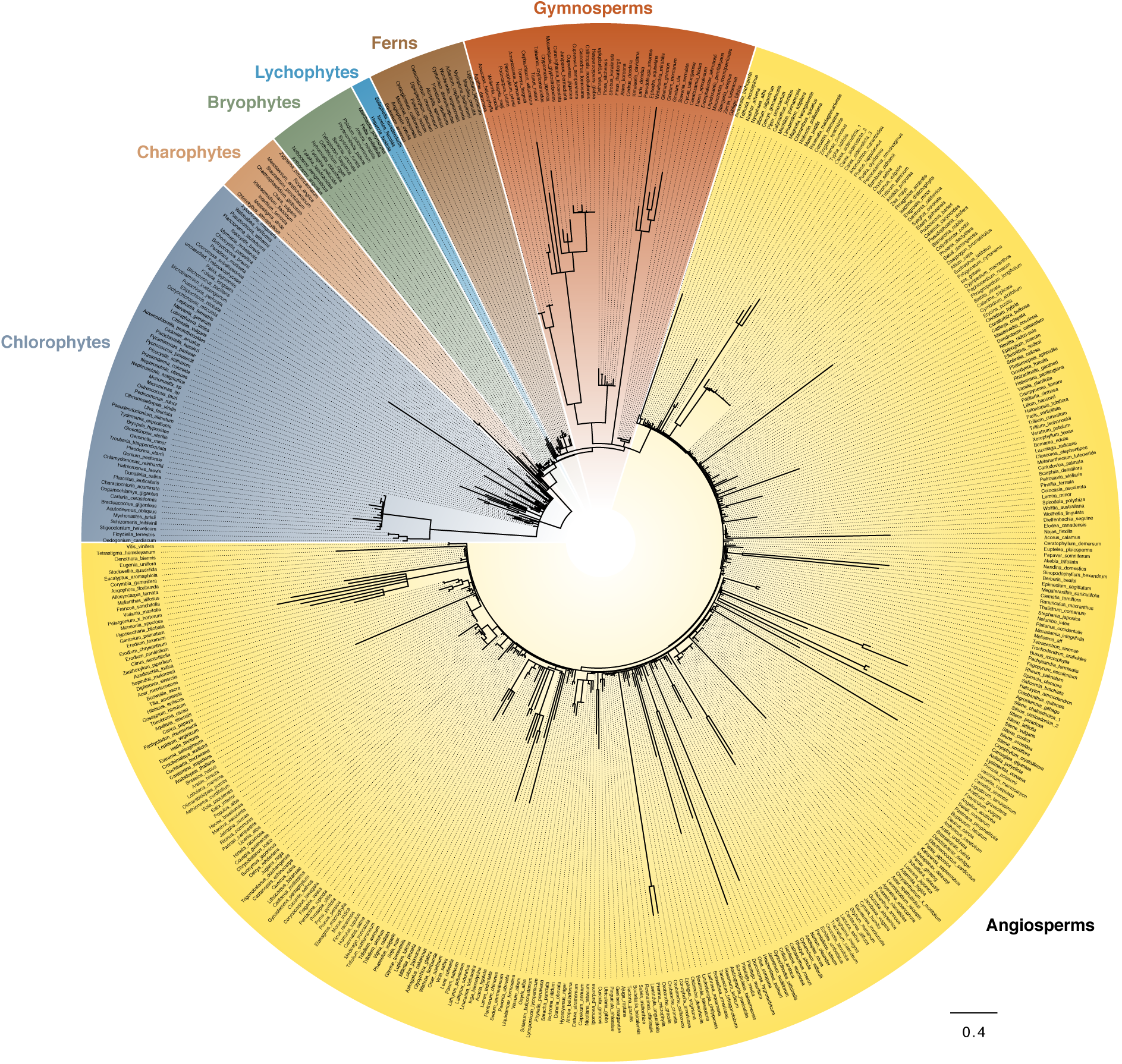
ClpP1 evolutionary rates across green plants. Tree is the same as shown in Figure 1 but includes species names for each branch. Branch length represents amino acid substitutions per site.

**Figure S2:**
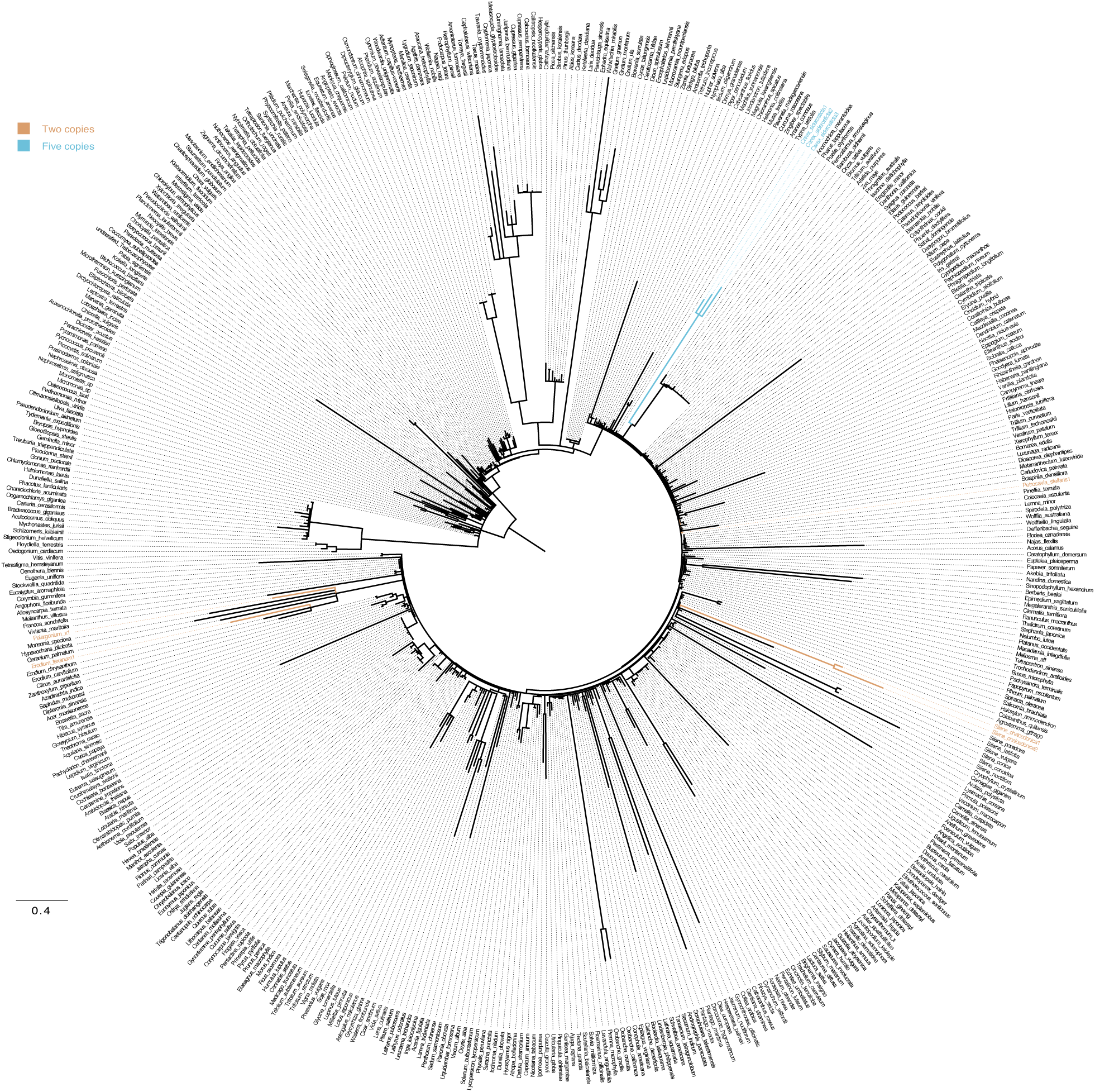
*clpP1* duplications across green plants superimposed on ClpP1 evolutionary rate tree. Internal branches colored based on simple parsimony.

**Figure S3:**
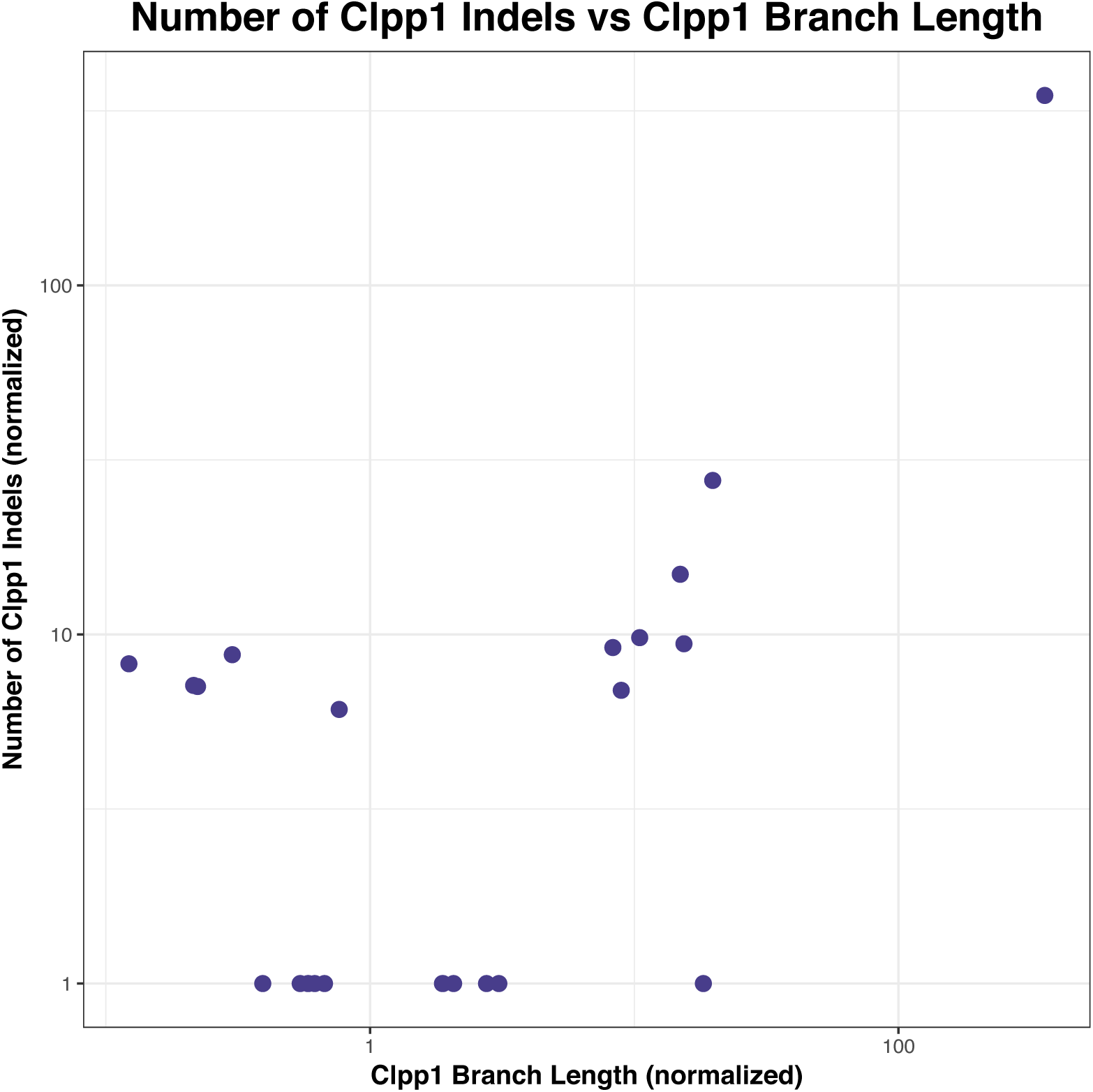
Correlation between *clpP1* indels and ClpP1 branch length. As with Figure 7, normalization of both axes was achieved using independent sets of non-Clp nuclear-encoded proteins (n=10, concatenated).

**Figure S4:**
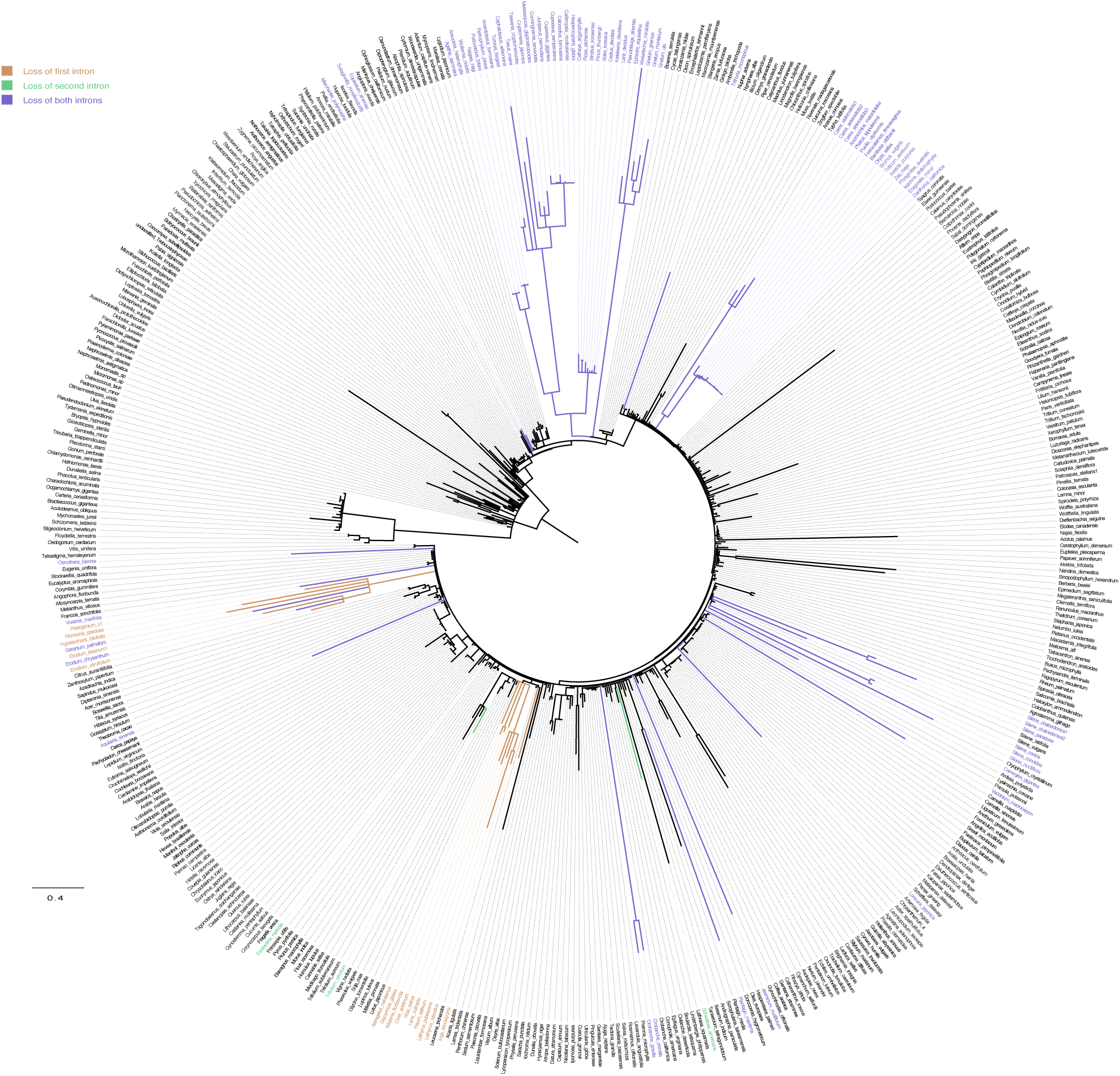
Loss of *clpP1* introns superimposed on ClpP1 evolutionary rate tree. Internal branches colored based on simple parsimony.

**Figure S5:**
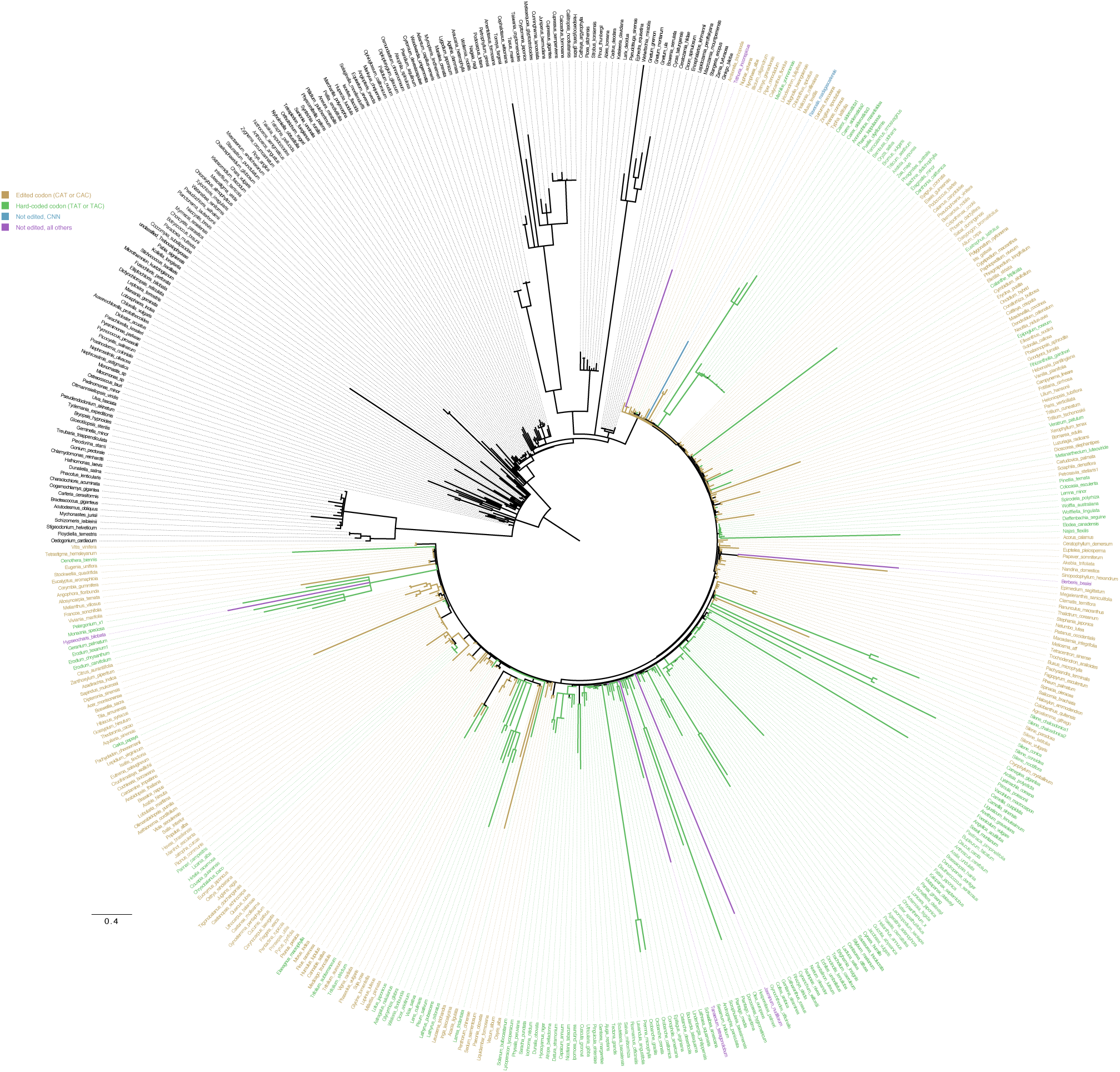
RNA editing of *clpP1* codon 187 superimposed on ClpP1 evolutionary rate tree. Internal branches colored based on simple parsimony.

**Figure S6:**
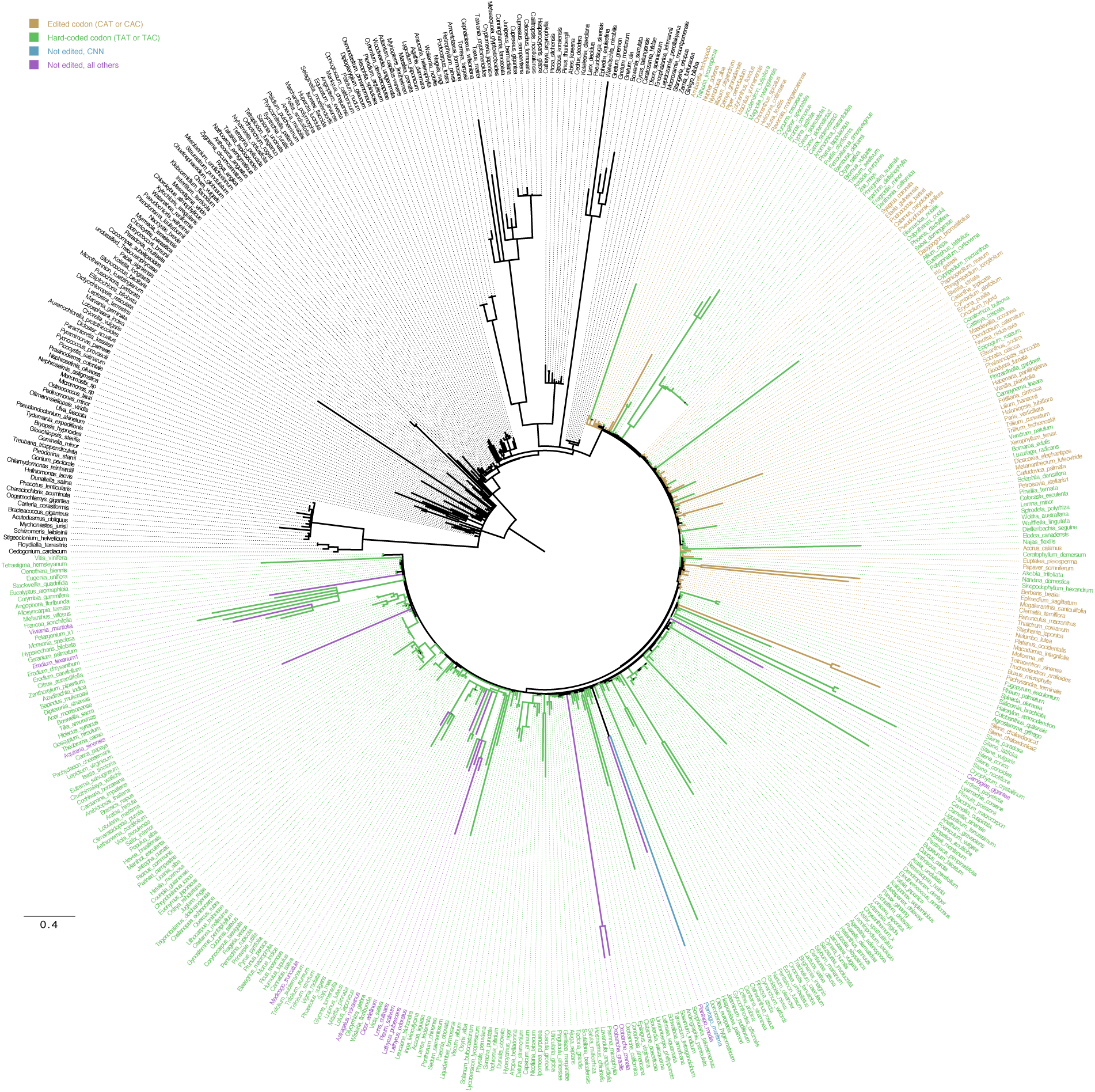
RNA editing of *clpP1* codon 28 superimposed on ClpP1 evolutionary rate tree. Internal branches colored based on simple parsimony.

**Figure S7:**
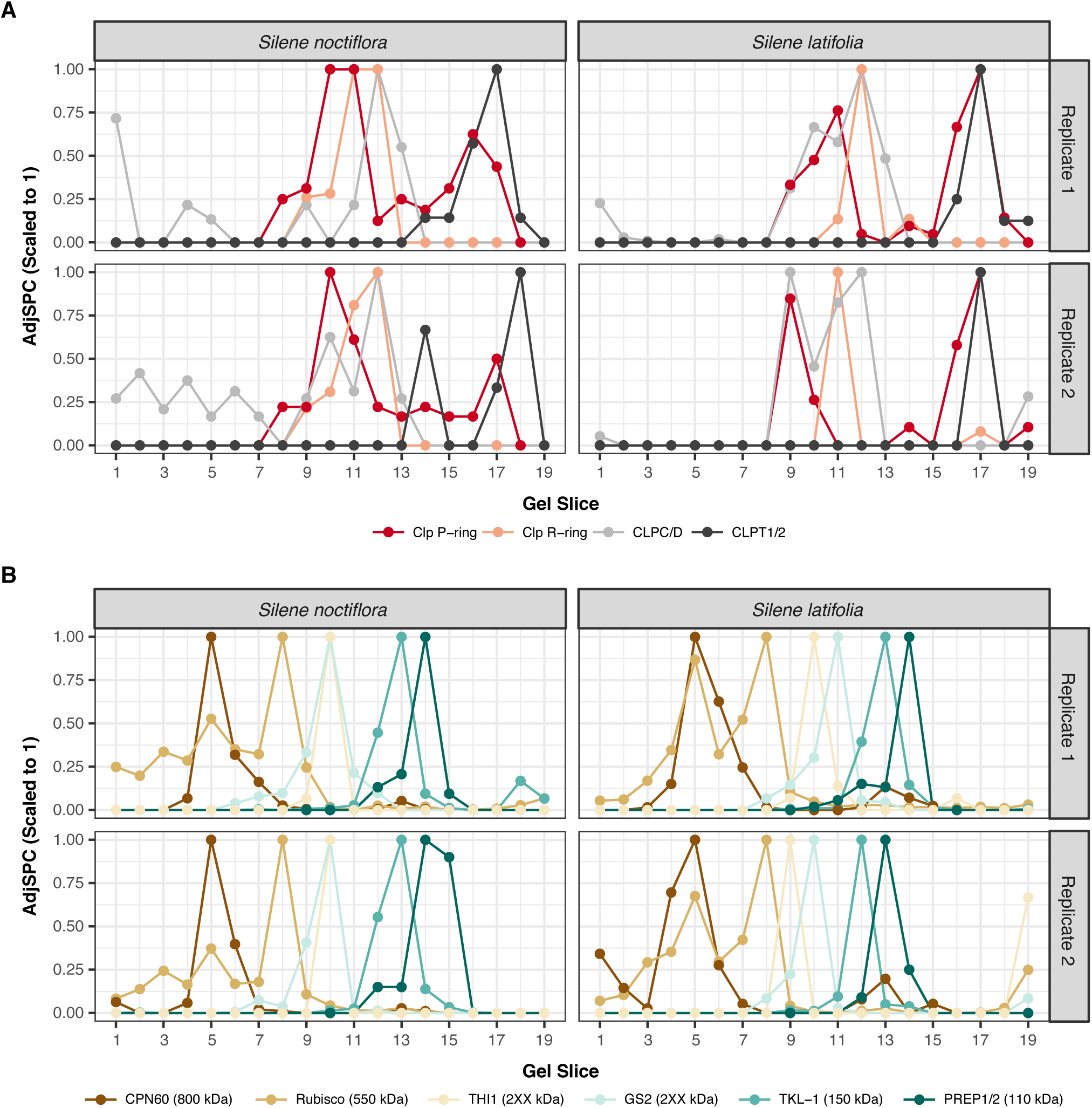
Size-fractionated mass spectrometry analysis of *Silene* plastid proteins. Gel slices correspond to native gel shown in Figure 3. A) Scaled AdjSPC summed across all subunits in different components of the Clp complex in *S*. *noctiflora* and *S*. *latifolia*. Values are scaled to 1 by dividing by the maximum observed value in any of the 19 gel slices. B) Control complexes used for internal calibration. Scaled AdjSPC values are calculated as in panel A.

**Figure S8:**
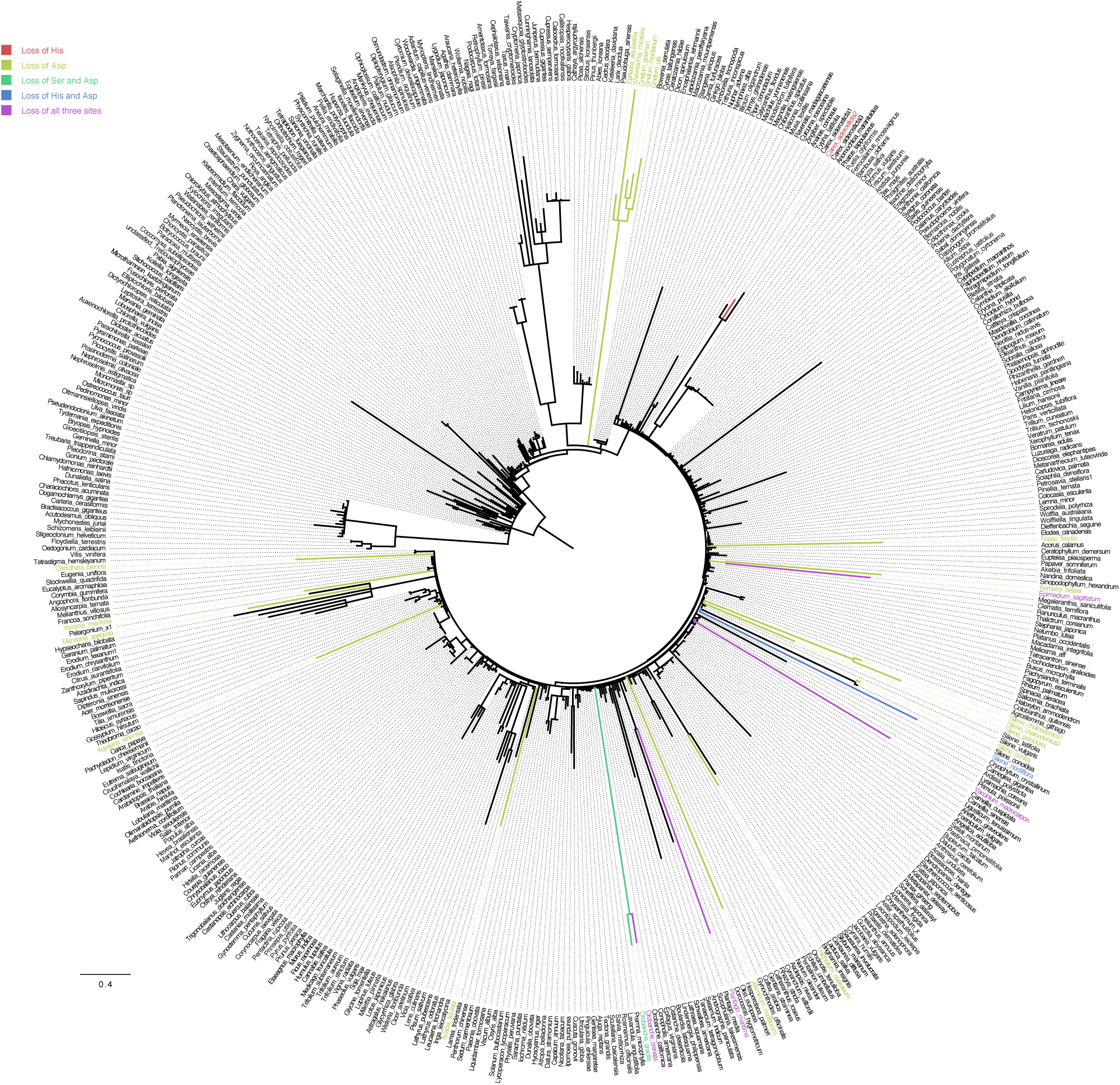
Loss of ClpP1 catalytic sites superimposed on ClpP1 evolutionary rate tree. Internal branches colored based on simple parsimony.

**Table S1.** *clpP1* and *psaA d*_N_, *d*_S_, and *d*_N_/*d*_S_ values in 25 angiosperms

| Species | *clpP1* $d_N$ | *clpP1* $d_S$ | *clpP1* $d_N/d_S$ | *psaA* $d_N$ | *psaA* $d_S$ | *psaA* $d_N/d_S$ |
| --- | --- | --- | --- | --- | --- | --- |
| Ancestor of *S. bicolor* and *O. sativa* | 0.149 | 0.292 | 0.511 | 0.009 | 0.147 | 0.058 |
| *Sorghum bicolor* | 0 | 0.061 | 0 | 0.003 | 0.058 | 0.046 |
| *Oryza sativa* | 0.012 | 0.038 | 0.323 | 0.007 | 0.073 | 0.096 |
| *Musa acuminata* | 0.019 | 0.077 | 0.252 | 0.004 | 0.084 | 0.047 |
| *Cucumis sativus* | 0.073 | 0.086 | 0.855 | 0.009 | 0.107 | 0.084 |
| *Prunus persica* | 0.031 | 0.087 | 0.357 | 0.001 | 0.085 | 0.015 |
| *Lathyrus sativus* | 0.211 | 0.285 | 0.742 | 0.002 | 0.07 | 0.031 |
| *Glycine max* | 0.021 | 0.184 | 0.111 | 0.011 | 0.199 | 0.055 |
| *Medicago truncatula* | 0.126 | 0.2 | 0.632 | 0.003 | 0.04 | 0.069 |
| *Acacia ligulata* | 0.386 | 0.263 | 1.468 | 0.002 | 0.033 | 0.054 |
| *Ricinus communis* | 0.009 | 0.156 | 0.057 | 0.004 | 0.076 | 0.048 |
| *Populus trichocarpa* | 0.019 | 0.064 | 0.301 | 0.001 | 0.074 | 0.016 |
| *Arabidopsis thaliana* | 0.044 | 0.158 | 0.277 | 0.006 | 0.147 | 0.041 |
| *Gossypium raimondii* | 0.015 | 0.069 | 0.214 | 0.005 | 0.1 | 0.048 |
| *Eucalyptus grandis* | 0.004 | 0.046 | 0.087 | 0.003 | 0.033 | 0.09 |
| *Oenothera biennis* | 0.365 | 0.478 | 0.764 | 0.002 | 0.108 | 0.022 |
| *Geranium maderense* | 0.493 | 0.424 | 1.161 | 0.009 | 0.169 | 0.05 |
| *Vitis vinifera* | 0.063 | 0.042 | 1.494 | 0.003 | 0.051 | 0.059 |
| *Erythranthe lutea* | 0.001 | 0.053 | 0.023 | 0.002 | 0.069 | 0.026 |
| *Plantago maritima* | 0.679 | 0.638 | 1.065 | 0.001 | 0.108 | 0.011 |
| *Solanum lycopersicum* | 0.13 | 0.13 | 0.999 | 0.002 | 0.093 | 0.026 |
| *Lobelia siphilitica* | 0.297 | 0.378 | 0.785 | 0.008 | 0.197 | 0.038 |
| *Silene noctiflora* | 0.578 | 0.532 | 1.085 | 0.003 | 0.019 | 0.134 |
| *Silene latifolia* | 0 | 0.047 | 0 | 0.001 | 0.004 | 0.136 |
| *Liriodendron chinense* | 0.008 | 0.039 | 0.199 | 0.003 | 0.047 | 0.061 |
| *Amborella trichopoda* | 0.034 | 0.135 | 0.249 | 0.006 | 0.135 | 0.046 |

**Table S2: Large dataset sampling**

**Table S3: Small dataset sampling**

**Dataset S1A: Mass spectrometry-based identification of *S*. *noctiflora* proteins and annotation based on best homolog to *Arabidopsis*.**

**Dataset S1B: Mass spectrometry-based identification of *S*. *latifolia* proteins and annotation based on best homolog to *Arabidopsis*.**

